# A time-delayed mechanochemical feedback model reconciles stable maintenance and dynamic remodeling of cell–matrix adhesions

**DOI:** 10.64898/2026.08.28.747716

**Authors:** Eiji Matsumoto, Sho Yokoyama, Tsubasa S. Matsui, Tom Araki, Shinji Deguchi

## Abstract

Focal adhesions maintain force-bearing attachment between cells and the extracellular matrix but can also undergo dynamic remodeling. Their assembly and actomyosin tension are coupled through mechanochemical feedback. The processes underlying this feedback are not instantaneous and therefore involve a time delay. However, how this delayed feedback gives rise to stable adhesion maintenance or dynamic remodeling remains unclear. Here, paired time-lapse measurements of vinculin fluorescence and traction stress revealed distinct local adhesion–force dynamics, including low-fluctuation and recurrent fluctuation patterns. To examine how these patterns could arise, we formulated a minimal mechanochemical model coupling focal adhesion assembly and actomyosin force through delayed reciprocal feedback. The model exhibited stable and oscillatory modes depending on feedback strength, the balance of opposing feedback effects, and the effective feedback delay. Bistability and hysteretic switching also occurred in a subset of parameter space, and the oscillation period followed a power-law relation with the delay. These results suggest that stable adhesion maintenance and dynamic remodeling can emerge from a common mechanochemical feedback architecture.

## I. INTRODUCTION

Adhesions between cells and the extracellular matrix (ECM) both transmit forces and dynamically assemble, remodel, and disassemble as cells spread, migrate, and respond to changes in their mechanical environment. Focal adhesions (FAs) are integrin-based mechanosensitive interfaces that connect the ECM to the actin cytoskeleton [1, 2]. Mechanosensing at FAs enables cells to detect local forces and the mechanical properties of the ECM and translate these cues into biochemical and cytoskeletal responses [3, 4]. For example, mechanical stress applied to integrins is transmitted through the cytoskeleton [5], and substrate stiffness regulates integrin–cytoskeleton coupling and FA formation and stability [6, 7]. Actomyosin (AM) contractility contributes to FA formation [8], and externally applied force can induce FA growth [9]. In turn, forces applied to integrins can activate intracellular signaling that promotes cytoskeletal and adhesion reinforcement [10]. FAs and the AM cytoskeleton thus form a reciprocally coupled mechanochemical system [11]. Although many molecular and mechanical interactions in this system have been characterized, how sustained force transmission and dynamic remodeling are coordinated within cell–ECM adhesions remains incompletely understood.

At the molecular level, tensile loading of talin exposes vinculin-binding sites, and talin–vinculin engagement reinforces integrin–cytoskeleton coupling during FA maturation [12, 13]. Signaling through integrins and FAs can regulate AM force in part through Rho-family GTPases [10]. RhoA signaling promotes myosin-II contractility and traction generation [8, 14]. Rac1-related signaling can promote actin-based protrusion and the turnover of small peripheral adhesions; it can also suppress RhoA signaling or limit myosin-II-dependent adhesion maturation [15–17]. These mechanisms suggest that reciprocal adhesion–force coupling is shaped by tension-dependent adhesion reinforcement and the balance between RhoAand Rac1-related effects on AM contractility.

Reciprocal adhesion–force feedback involves signaling, cytoskeletal reorganization, force generation, and adhesion remodeling and therefore does not operate instantaneously. Time-resolved measurements at individual FAs have revealed temporal offsets between adhesion assembly and FAK activation [18] and between traction and FAK activation [19]. Together, these finite response times can introduce an effective feedback delay, altering system stability and giving rise to complex temporal behavior [20, 21]. Despite this potential dynamical importance, mathematical models of cell–ECM adhesion dynamics rarely include explicit feedback delays. Previous models have coupled FA formation to stress-fiber assembly and contractile tension [22, 23] or linked bidirectional adhesion signaling and AM mechanics to intermittent leading-edge motility [24]. A mechanochemical model predicted bistability and hysteresis in Rho-dependent cellular contractility [25]. Traction oscillations have been generated through motor–clutch loading [26] and actin-network feedback [27]. However, it remains unclear how a single mechanochemical feedback architecture, in which FA assembly and AM tensile force are reciprocally coupled with an effective delay, can give rise to both stable adhesion maintenance and dynamic adhesion remodeling.

Here, we analyzed paired time-lapse measurements of local RFP–vinculin fluorescence and traction stress obtained by wrinkle force microscopy in A7r5 vascular smooth muscle cells [28]. The amplitudes of temporal fluctuations in both signals varied across space, and the local time courses included low-fluctuation and recurrent fluctuation patterns. To explore how these patterns could arise, we formulated a minimal model of delayed reciprocal feedback between FA assembly and AM tensile force. The model linked stable and oscillatory dynamics to feedback strength, balance, and delay, suggesting a common mechanochemical basis for stable adhesion maintenance and dynamic remodeling.

## II. RESULTS

### A. Local cell–ECM adhesion dynamics exhibit low-fluctuation and recurrent fluctuation patterns

We first examined temporal variation in local cell–ECM adhesion dynamics using paired time-lapse measurements of RFP–vinculin fluorescence and traction stress (Fig. 1A and Supplementary Movie S1). We analyzed the local time series to compare fluctuation magnitudes and temporal patterns in the two signals (Fig. 1B–G). The local time series included both low-fluctuation and recurrent fluctuation patterns. In the low-fluctuation example, the smoothed RFP–vinculin and traction signals showed comparatively small relative deviations, and the joint trajectory occupied a compact region (Fig. 1B–D). By contrast, the recurrent fluctuation example showed increases and decreases in both signals, and the corresponding joint trajectory followed a broad, loop-shaped path in the RFP–traction plane (Fig. 1E–G). Additional examples from each time-lapse dataset exhibited low-fluctuation or recurrent fluctuation patterns (Fig. S2). The two representative patterns were qualitatively preserved across the examined smoothing windows (Fig. S3).

**FIG. 1.**
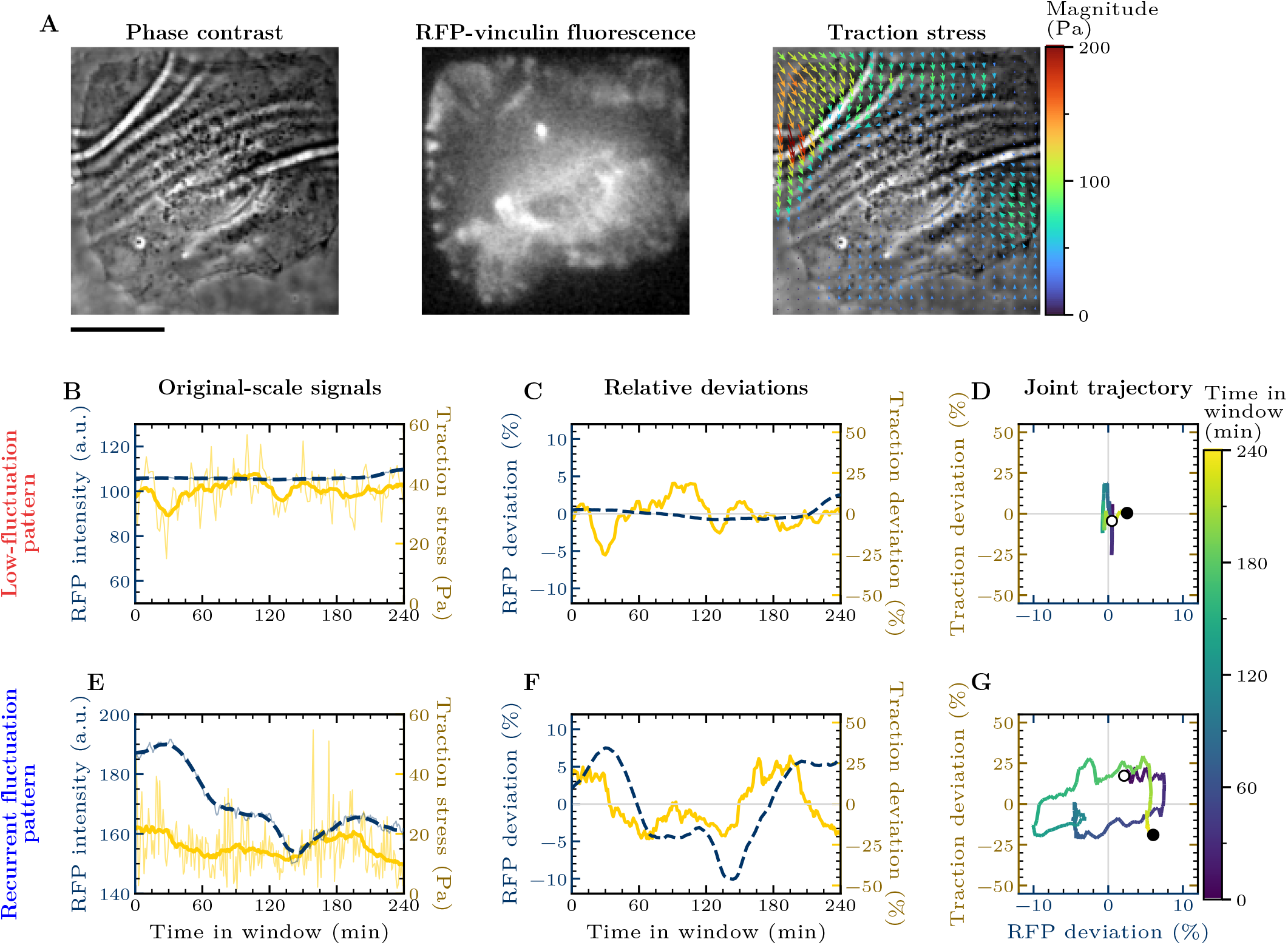
Local adhesion and traction signals exhibit heterogeneous temporal dynamics. Representative low-fluctuation and recurrent fluctuation patterns differ in fluctuation magnitude and joint-trajectory geometry. **(A)** Representative phase-contrast and RFP–vinculin images are shown with traction-stress vectors overlaid on phase contrast; the corresponding time-lapse sequence is provided in Supplementary Movie S1. Vector orientation indicates traction direction, and vector length and color encode traction magnitude. The scale bar represents 20 *µ*m. **(B–D)** The low-fluctuation example shows relatively limited deviations and a compact joint trajectory. **(E–G)** The recurrent fluctuation example shows larger fluctuations and a broad, loop-shaped joint trajectory. In panels B and E, thin blue and yellow lines show the raw RFP–vinculin and traction signals, respectively, and the corresponding thick lines show the smoothed signals. In panels C and F, relative deviations were obtained by linearly detrending each smoothed signal within the displayed 240-min window and expressing the residual as a percentage of the corresponding recording-wide median signal level. In panels D and G, the relative RFP and traction deviations are plotted as joint trajectories colored by time. Open and filled circles mark the first and final points, respectively.

### B. Amplitudes of adhesion and traction fluctuations vary across space and recordings

We next quantified local fluctuation amplitudes of RFP–vinculin fluorescence and traction stress across the time-lapse datasets (Fig. S1). Both amplitude measures varied across grid positions and among the datasets (Fig. 2A and B). The corresponding density estimates differed in location and shape among the datasets, while the equally weighted aggregate for each signal spanned a broad amplitude range (Fig. 2C and D). Alternative normalization and distribution analyses yielded similarly broad amplitude ranges without robust evidence for discrete amplitude populations (Fig. S4 and Fig. S5). These observations motivated us to examine whether a common mechanochemical feedback architecture could generate both stable and dynamic adhesion behaviors.

**FIG. 2.**
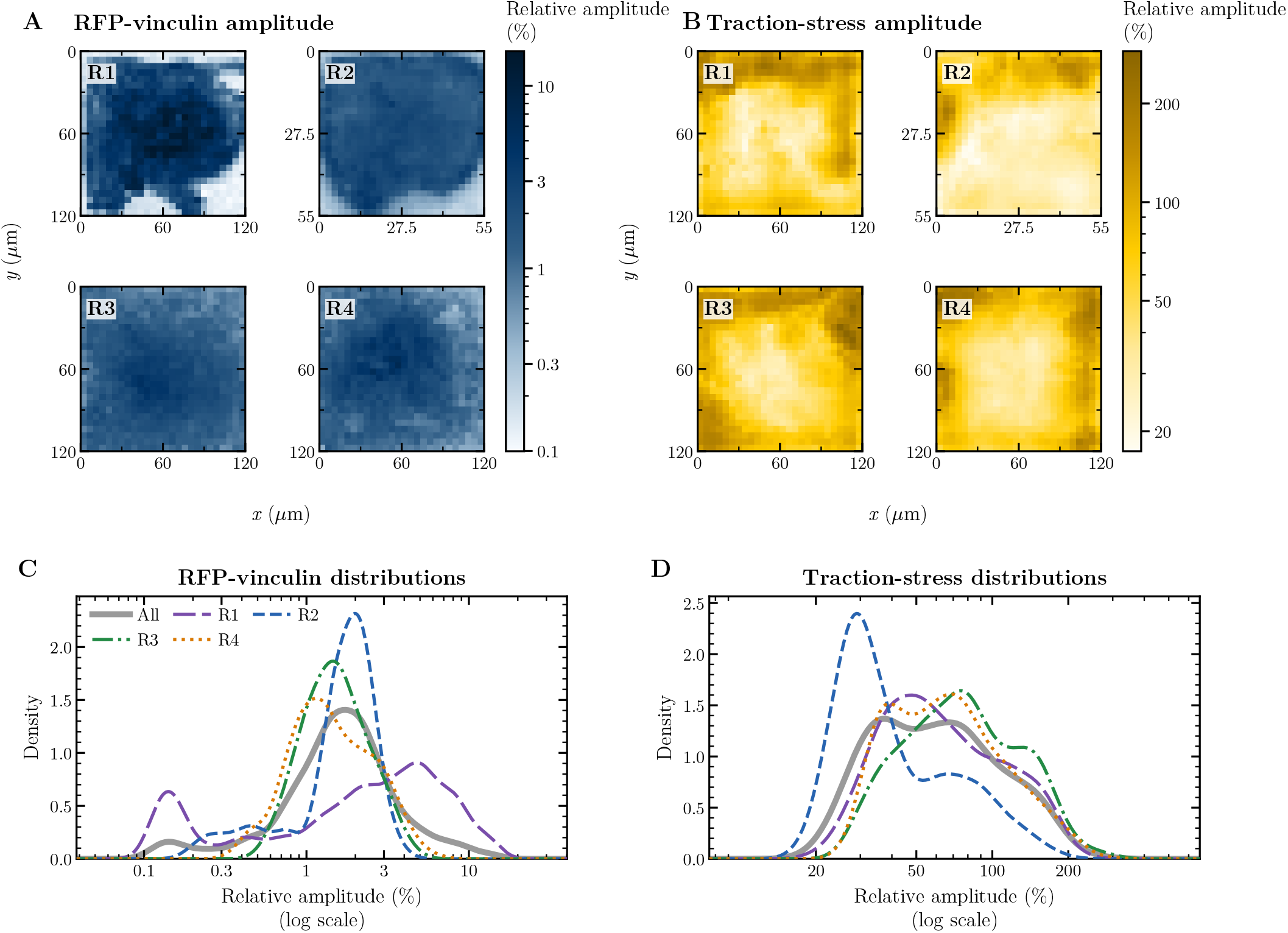
Local adhesion and traction signals exhibit heterogeneous fluctuation amplitudes. Spatial maps and density distributions reveal variation in the amplitudes of both signals across the measured regions. **(A and B)** Spatial maps show relative RFP–vinculin and traction-stress amplitudes, respectively, for the four time-lapse datasets (R1–R4). Each map preserves the physical dimensions of the corresponding analyzed region. Within each window, relative amplitude was calculated from the 10th-to-90th percentile range of the linearly detrended signal and normalized by the corresponding dataset-wide median signal level. For each spatial position, the median relative amplitude across overlapping 240-min windows shifted in 30-min steps is shown as a percentage. The color bars use logarithmic scales. **(C and D)** Kernel-density estimates show the corresponding RFP–vinculin and traction-stress amplitude distributions, respectively, for R1–R4 and their equally weighted aggregate (All).

### C. A time-delayed mechanochemical feedback model generates stable and oscillatory adhesion dynamics

We constructed a dimensionless model coupling focal adhesion assembly, *A*(*t*), and actomyosin tensile force, *F* (*t*), through reciprocal delayed feedback (Fig. 3A). Their dynamics are governed by

**FIG. 3.**
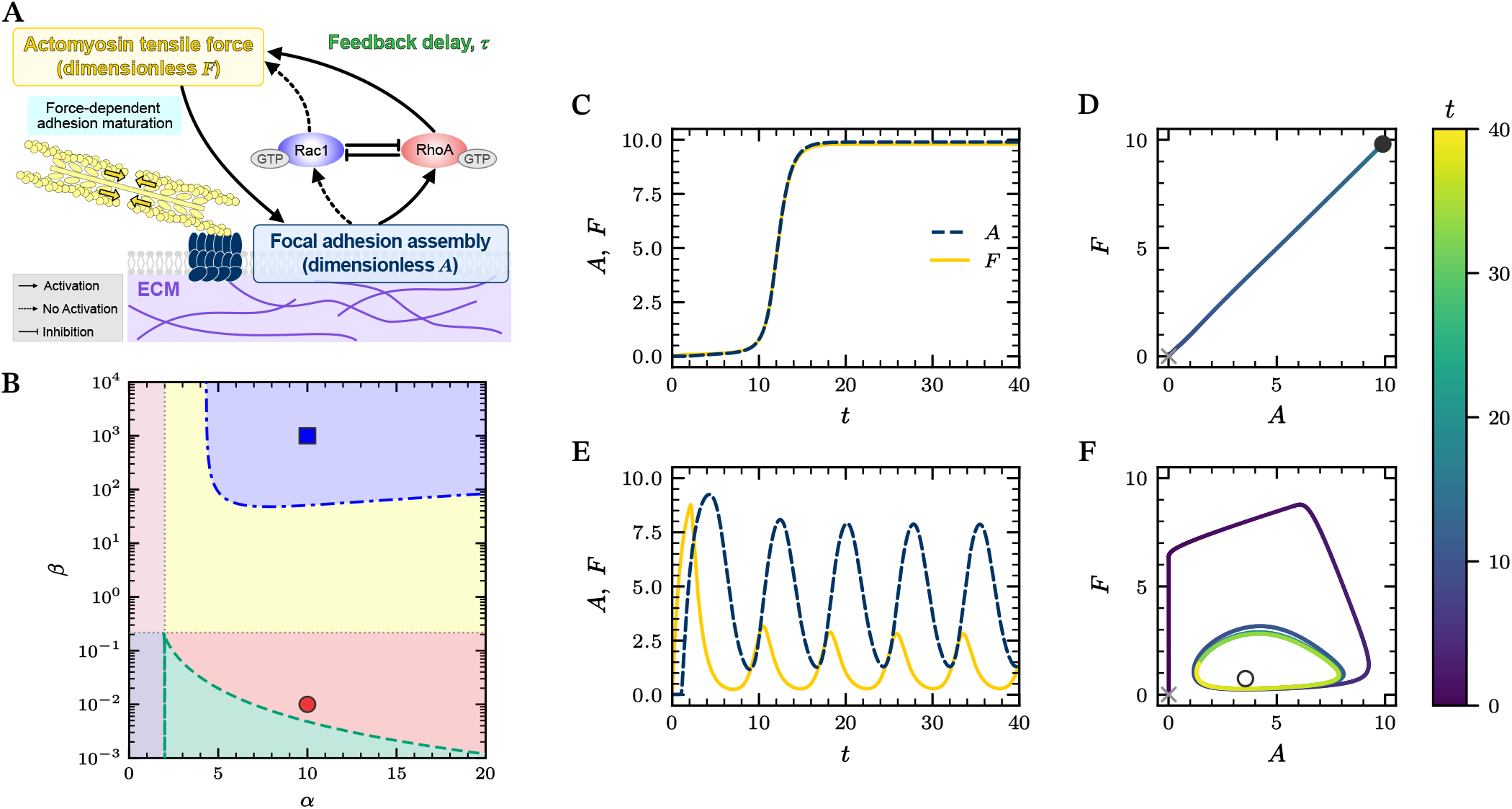
Delayed mechanochemical feedback generates stable fixed-point behavior and sustained limit-cycle oscillations. The phase diagram and representative trajectories show that the same focal adhesion–actomyosin feedback architecture produces either a stable fixed point or a sustained limit cycle under different parameter conditions. **(A)** The dimensionless variables *A* and *F* represent focal adhesion assembly and actomyosin tensile force, respectively. The schematic shows force-dependent adhesion assembly and maturation together with the opposing RhoA-related activating and Rac1-related inhibitory effects of adhesion assembly on actomyosin tensile force. The feedback interactions involve multistep molecular cascades whose cumulative delay is represented in the model by a single effective feedback delay, *τ* . **(B)** A representative phase diagram shows the model’s dynamical organization in the (*α, β*) plane at *τ* = 1. Colors distinguish inactive (purple), low-activity stable (pink), high-activity stable (red), and adhesion-dominant (yellow) subdivisions of monostable fixed-point behavior, together with bistable (green) and oscillatory (blue) regimes. Green dashed and blue dash-dotted curves denote saddle-node and Hopf bifurcation boundaries, respectively. **(C and D)** At (*α, β, τ* ) = (10, 0.01, 1), the time courses converge to a high-activity stable fixed point (C), and the phase-plane trajectory approaches the same fixed point (D). **(E and F)** At (*α, β, τ* ) = (10, 1000, 1), the time courses exhibit sustained oscillations (E), and the phase-plane trajectory approaches a stable limit cycle surrounding an unstable fixed point (F). In panels C and E, *A*(*t*) is shown by the navy dashed line and *F* (*t*) by the yellow solid line. In panels D and F, trajectory color indicates time, gray crosses mark the constant history values (*A*_0_, *F*_0_) = (0.01, 0.01) prescribed for *t* ∈ [−*τ*, 0], and filled and open circles denote stable and unstable fixed points, respectively.

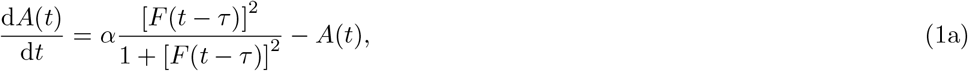

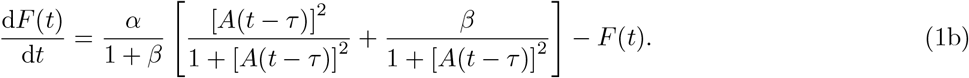

Here, *α* is the mechanochemical coupling strength, *β* is the ratio of Rac1-related inhibitory to RhoA-related activating feedback strengths, and *τ* is the effective feedback delay. AM tensile force promotes FA assembly through tension-dependent FA maturation (Eq. (1a)). FA assembly feeds back through two signaling arms, with RhoA-related activation promoting AM tensile force and Rac1-related inhibition opposing it (Eq. (1b)). These feedback loops involve multistep molecular and structural processes that introduce finite response delays, represented collectively in the model by the single effective feedback delay *τ* . The model formulation and analysis procedures are described in detail in the Materials and Methods, with the derivation and nondimensionalization provided in Supplementary Text S1.

Using *τ* = 1 as a representative feedback delay, we mapped the model’s dynamical organization in the (*α, β*) plane and identified four descriptive subdivisions of monostable fixed-point behavior, as well as a bistable regime and an oscillatory regime (Fig. 3B). The four monostable subdivisions were defined by the absolute and relative fixed-point levels of *A*^∗^ and *F* ^∗^ (Fig. S8). At (*α, β, τ* ) = (10, 0.01, 1), the time courses of *A*(*t*) and *F* (*t*) converged to constant values, and the phase-plane trajectory approached a stable fixed point, providing a representative example of the high-activity stable mode (Fig. 3C and D). By contrast, at (*α, β, τ* ) = (10, 1000, 1), both variables exhibited sustained oscillations, and the phase-plane trajectory approached a stable limit cycle surrounding an unstable fixed point, providing a representative example of the oscillatory mode (Fig. 3E and F). The same delayed mechanochemical feedback architecture therefore generated either stable fixed-point or oscillatory limit-cycle dynamics under different parameter conditions. For fixed *α* and *τ*, the relative balance between the two feedback effects, set by *β*, determined which mode emerged.

### D. Feedback strength, balance, and delay organize distinct dynamical regimes

We mapped phase diagrams in the (*α, β*) plane at *τ* = 0, 0.5, 1, and 5 to examine how feedback strength (*α*), feedback balance (*β*), and delay (*τ* ) affected the dynamical regimes and the occurrence of bistability and hysteresis (Fig. 4A). At *τ* = 0, the parameter plane contained monostable fixed-point and bistable regimes but no oscillatory regime. For nonzero feedback delays (*τ >* 0), the fixed point lost stability across a Hopf bifurcation boundary, with sustained oscillations observed on its oscillatory side. The oscillatory region expanded as the boundary shifted with increasing *τ* . Across these delay conditions, variations in *α* and *β* organized the inactive, low-activity stable, high-activity stable, adhesion-dominant, bistable, and oscillatory regions.

**FIG. 4.**
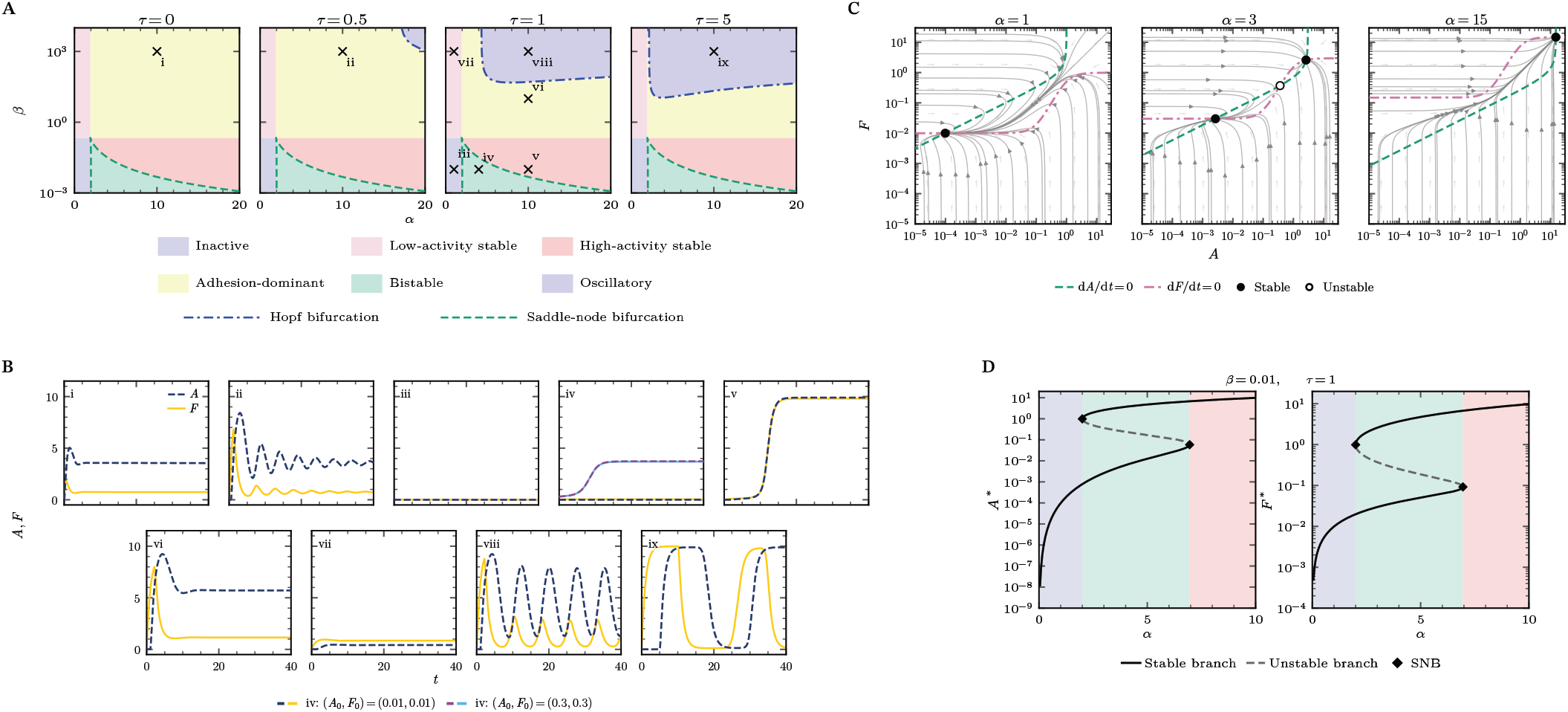
Feedback parameters and delay reorganize the model’s dynamical landscape. Feedback strength, balance, and delay select stable, oscillatory, or bistable responses, and the bistable branch structure permits hysteresis. **(A)** Phase diagrams in the (*α, β*) plane at *τ* = 0, 0.5, 1, and 5 show no oscillatory regime at *τ* = 0. For nonzero feedback delays (*τ >* 0), a Hopf bifurcation boundary separates stable fixed-point and oscillatory regions, and the oscillatory region expands as the boundary shifts with increasing *τ* . Colors distinguish inactive, low-activity stable, high-activity stable, adhesion-dominant, bistable, and oscillatory regions. Blue dash-dotted and green dashed curves denote Hopf and saddle-node bifurcation boundaries, respectively. Markers i–ix indicate the conditions examined in panel B. **(B)** Time courses i–ix illustrate adhesion-dominant (i, ii, and vi), inactive (iii), bistable (iv), high-activity stable (v), low-activity stable (vii), and oscillatory (viii and ix) responses. All cases use the constant history (*A*_0_, *F*_0_) = (0.01, 0.01). Bistable case iv additionally shows the alternative history (0.3, 0.3), plotted as indicated in the legend, which converges to a different stable fixed point. **(C)** For the nondelayed ordinary differential equation (ODE) system (*τ* = 0) at *β* = 0.01, phase planes show an inactive stable fixed point at *α* = 1, bistability at *α* = 3, and a high-activity stable fixed point at *α* = 15. Green dashed and pink dash-dotted curves denote the d*A/*d*t* = 0 and d*F/*d*t* = 0 nullclines, respectively, and gray streamlines show the corresponding ODE vector field. **(D)** Steady-state values *A*^*∗*^ and *F*^*∗*^ are plotted against *α* at *β* = 0.01 and *τ* = 1, revealing the bistable branch structure that permits hysteresis under quasistatic variation of *α*. Solid black and dashed gray curves denote stable and unstable branches, respectively. Black diamonds mark saddle-node bifurcation (SNB) points, and the background colors indicate the inactive, bistable, and high-activity stable intervals corresponding to the *τ* = 1 phase diagram in panel A.

We then compared the temporal responses at the parameter conditions marked i–ix in the phase diagrams (Fig. 4B). Case iii converged to an inactive fixed point with both *A* and *F* near zero, case vii to a low-activity stable fixed point with both variables remaining low but finite, and case v to a high-activity stable fixed point with both variables approaching high levels. Cases i, ii, and vi converged to adhesion-dominant fixed points with higher steady-state levels of *A* than *F* . Cases i and vi approached these fixed points after a pronounced overshoot, whereas case ii converged to its fixed point through damped oscillations. Comparison of cases i, ii, viii, and ix at the common parameter condition (*α, β*) = (10, 1000) showed that increasing *τ* transformed the convergent responses at *τ* = 0 and 0.5 into sustained oscillations at *τ* = 1 and 5. Compared with *τ* = 1, the oscillations at *τ* = 5 had larger amplitudes and a longer period. In bistable case iv, the constant histories (*A*_0_, *F*_0_) = (0.01, 0.01) and (0.3, 0.3) converged to inactive and high-activity stable fixed points, respectively, under the same parameter condition. Parameter values affected both transient and long-term behavior, and within the bistable regime the final state depended on the system’s history.

To determine how *α* altered the number and stability of fixed points and their basins of attraction, we examined the nullclines and phase-plane flow of the nondelayed model (*τ* = 0) at *β* = 0.01 (Fig. 4C). At *α* = 1 and 15, the nullclines intersected at a single inactive stable fixed point and a single high-activity stable fixed point, respectively, and the streamlines converged toward the corresponding attractors. At *α* = 3, the nullclines intersected at two stable fixed points with an unstable fixed point between them, and the streamlines separated into distinct basins leading to the inactive and high-activity stable attractors. Extended fixed-point maps and phase portraits further illustrated these monostable and bistable structures across parameter conditions (Fig. S8). At *β* = 0.01 and *τ* = 1, saddle-node bifurcations at *α* ≈2.00 and 6.95 bounded a bistable interval (Fig. 4D). Within this interval, lower and upper stable steady states coexisted, with an unstable branch lying between them. Under quasistatic variation of *α*, the state would jump from the lower to the upper stable branch at the upper saddle-node during an increasing sweep and from the upper to the lower stable branch at the lower saddle-node during a decreasing sweep. The different thresholds for increasing and decreasing sweeps produced hysteresis.

### E. Oscillation period scales with feedback delay, phase difference remains near *π/*2, and parameters jointly determine amplitudes

We next examined how feedback strength *α*, feedback balance *β*, and delay *τ* affected the period, phase difference, and amplitudes of the oscillatory mode. We found that the oscillation period *T* scaled with feedback delay *τ* according to the power-law relation *T* = 6.88*τ* ^0.854^ (*R*^2^ = 0.995; Fig. 5A). Separate power-law fits for fixed values of *α* or *β* yielded similar exponents and consistently high *R*^2^ values, indicating that the delay–period relationship was consistent across the examined feedback parameter values (Fig. S6).

**FIG. 5.**
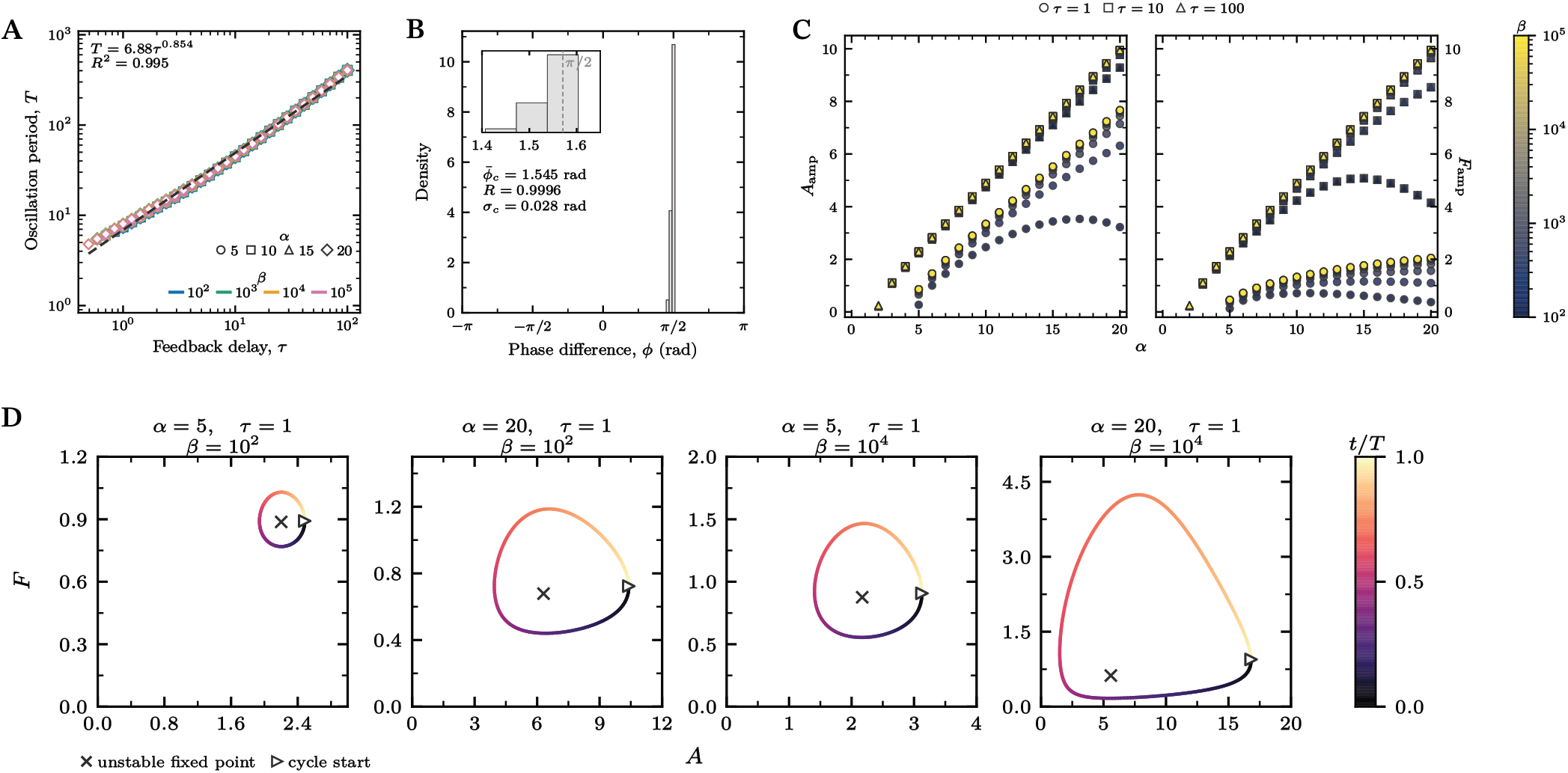
Oscillation period scales with feedback delay, phase difference remains near *π/*2, and feedback parameters jointly shape oscillation amplitudes. Period and phase are quantified across the oscillatory region, while amplitude measurements and representative cycles show how feedback parameters alter cycle size and geometry. **(A)** Oscillation period *T* is plotted against feedback delay *τ* for selected parameter combinations, with marker shape and color denoting *α* and *β*, respectively. The black dashed line shows the fitted power-law scaling relation *T* = 6.88*τ* ^0.854^, with coefficient of determination *R*^2^ = 0.995. **(B)** The density distribution shows the circular phase difference *ϕ* across oscillatory parameter combinations, with positive values indicating that *F* leads *A*. The inset enlarges the distribution around its circular mean, 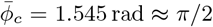, with mean resultant length *R* = 0.9996 and circular standard deviation *σ*_*c*_ = 0.028 rad; the gray dashed line marks *π/*2. **(C)** Half-range amplitudes *A*_amp_ and *F*_amp_, calculated as half the difference between the mean post-transient peak value and the mean post-transient trough value, are plotted against *α*, with marker shape and color denoting *τ* and *β*, respectively. **(D)** Representative oscillatory trajectories over one post-transient cycle are shown for the indicated parameter combinations, with color indicating normalized cycle time *t/T* .

We then quantified the circular phase difference *ϕ* between *A* and *F*, with positive values indicating that *F* led *A* (Fig. 5B). At the first Hopf boundary, the critical linear mode has an exact phase difference of *π/*2, with *F* leading *A* (Eq. (S51)). Across the finite-amplitude oscillatory solutions, the numerically estimated phase difference remained tightly distributed near this value, with a circular mean of 1.545 rad, a mean resultant length of *R* = 0.9996, and a circular standard deviation of 0.028 rad. Parameter-resolved circular means also remained near *π/*2 across the examined values of *α* and *β* (Fig. S7).

We next examined how *α, β*, and *τ* shaped the half-range amplitudes *A*_amp_ and *F*_amp_. Across most of the oscillatory region, *A*_amp_ increased approximately linearly with *α*, with the fitted slope approaching 0.5 as *τ* increased (Fig. 5C and Fig. S7A). *F*_amp_ also generally increased with *α*, but its *α* dependence remained weaker at short delays or low *β* (Fig. S7A and B). At fixed *α* and *β*, increasing *τ* increased both amplitudes (Fig. 5C). At fixed *α* and *τ*, increasing *β* had a larger relative effect on *F*_amp_ than on *A*_amp_ (Fig. 5C and Fig. S7C). Consequently, at larger *τ* and *β*, both amplitudes approached the common upper envelope *α/*2, and their ratio approached unity.

Representative limit cycles at *τ* = 1 illustrated the corresponding changes in cycle size and geometry (Fig. 5D). At the lower displayed values of *α* and *β*, the cycle remained compact; increasing *α* expanded its extent primarily along**RhoA**

*A*, whereas increasing *β* increased its extent along *F* relative to *A*. Despite differences in cycle size and geometry, the normalized-time color progression followed a similar pattern across the displayed trajectories, consistent with the near-constant phase relationship. The oscillation characteristics showed distinct parameter dependencies. The period scaled primarily with *τ*, the phase difference varied little around *π/*2, and the amplitudes changed gradually, with *α* setting their common upper envelope and *β* and *τ* controlling their approach to this envelope and their relative balance.

## III. DISCUSSION

Cell–ECM adhesions transmit contractile forces while continually undergoing molecular and structural remodeling. Although many of the underlying molecular interactions have been identified, how they collectively give rise to distinct stable and dynamic adhesion behaviors remains incompletely understood. We observed temporal and spatial variation in local RFP–vinculin fluorescence and traction stress, including low-fluctuation and recurrent fluctuation patterns. To examine how these temporal behaviors could arise, we formulated a time-delayed mechanochemical feedback model that reciprocally couples FA assembly and AM tensile force. For fixed feedback strength and delay, the model generated a high-activity stable mode or an oscillatory mode depending on the balance between the opposing RhoA- and Rac1-related adhesion-to-force feedbacks. These distinct dynamical modes provide a mechanochemical interpretation of stable adhesion maintenance and dynamic adhesion remodeling as distinct outcomes of a common mechanochemical feedback architecture (Fig. 6).

**FIG. 6.**
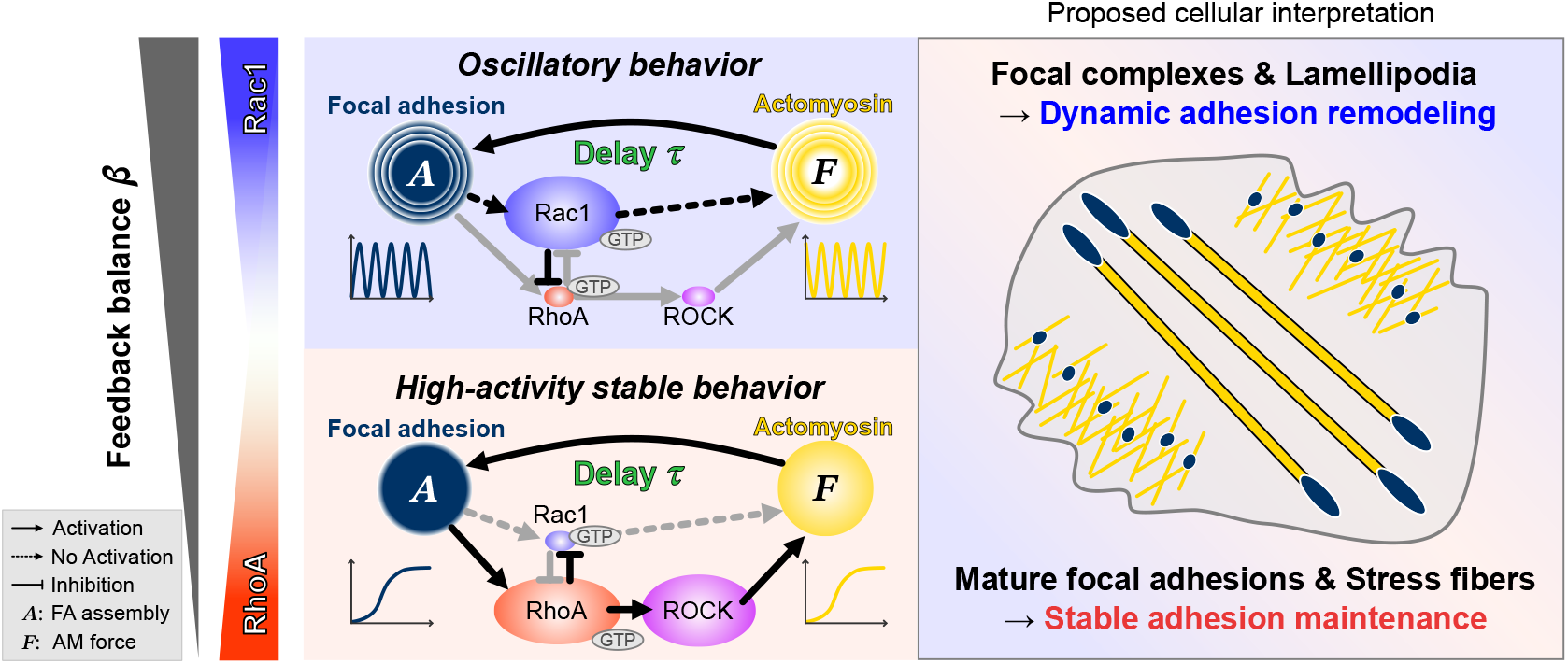
A mechanochemical interpretation of stable and oscillatory behaviors in cell–ECM adhesion. The schematic summarizes the feedback-dependent emergence of these behaviors and their proposed cellular interpretation. In the delayed feedback model, *β* sets the balance between Rac1-related inhibitory and RhoA-related activating feedbacks and, together with *α* and *τ*, determines whether the system exhibits high-activity stable or oscillatory behavior. High-activity stable behavior may correspond to adhesion maintenance involving mature FAs and stress fibers, and oscillatory behavior to dynamic remodeling involving focal complexes and lamellipodia.

This cellular interpretation is consistent with findings from previous experimental studies manipulating RhoA- and Rac1-related signaling. Microinjection and live-cell perturbations showed that Rac favored small peripheral focal complexes associated with cell-front protrusion, whereas Rho promoted their maturation into larger FAs [15, 29]. In cells with independently inducible constitutively active RhoA and Rac1, RhoA activation suppressed spreading and motility, Rac1 activation promoted dynamic protrusions and migration, and the two phenotypes could be switched reversibly [30]. Persistent Rho activity was associated with increased cell–ECM adhesion, reduced FA turnover, and impaired migration [31, 32]. Conversely, perturbations of the Rac1-related *β*-PIX–PAK1–paxillin module shifted adhesions toward either smaller, more dynamic or larger, longer-lived structures [17, 33]. Cell-generated traction was closely associated with FA assembly [34], and externally applied force promoted FA growth [9]. Integrin loading activated RhoA [10], contractility inhibition reduced adhesion strengthening or stability [35, 36], and optogenetic RhoA activation increased AM recruitment and traction [14, 37]. Across experimental systems and spatiotemporal scales, traction measurements, alone or combined with Förster resonance energy transfer (FRET) imaging, have identified stable, fluctuating, and oscillatory patterns of adhesion-associated force transmission [19, 38]. The modeled stable and oscillatory modes should therefore be interpreted as different balances between RhoA- and Rac1-related feedbacks rather than as states controlled exclusively by either pathway. Our model and these experimental findings suggest that this balance can bias cell–ECM adhesion dynamics toward stable maintenance with sustained force transmission or dynamic remodeling with force fluctuations (Fig. 6).

Whereas *β* sets the relative feedback balance, *α* represents a coarse-grained measure of the overall strength of the bidirectional mechanochemical coupling between FA assembly and AM tensile force. Thus, variation in *α* may reflect the combined influence of the extracellular mechanical and adhesive environment, adhesion-mediated force transmission, and cellular contractile activity. Experimentally, stiffer substrates promote myosin-II activity and traction generation [39] and are associated with more stable FAs [7]. Closer integrin–ligand spacing also promotes stable FA formation [40]. Integrin composition regulates myosin-II activation in response to the mechanical environment [41], force-sensitive engagement of talin and vinculin reinforces coupling between integrins and the actin cytoskeleton [12, 42], and myosin-II activity supports force generation and FA maturation [43, 44]. By integrating these effects into *α*, the model suggests that perturbations acting through different mechanisms can lead to similar adhesion dynamics when they produce comparable changes in overall adhesion–force coupling.

Although the descriptive subdivisions of monostable fixed-point behavior identified in our analysis do not map one-to-one onto molecular adhesion classes, they can be interpreted as qualitatively distinct states of cell–ECM adhesion. The inactive, low-activity stable, and high-activity stable modes may reflect, respectively, failure to establish persistent force-bearing adhesions, a weak adhesion state dominated by nascent adhesions or small focal complexes, and mature FAs associated with sustained AM tension. In particular, the high-activity stable mode is consistent with large, stable supermature FAs in contractile myofibroblasts [45], stronger and less dynamic traction with longer-lived FAs in stationary cells [46], and reduced traction fluctuations at FAs of vascular smooth muscle cells beyond a threshold substrate displacement [47]. The adhesion-dominant mode instead describes relatively high adhesion assembly compared with tensile force, consistent with observations that mature FA morphology and traction need not remain coupled [48]. Fibrillar adhesions may share some qualitative features of this mode [49, 50], although detailed molecular differences among adhesion types are outside the scope of the model. These subdivisions therefore provide a coarse-grained interpretation of stable adhesion states in terms of overall activity and adhesion–force balance.

In our model, a nonzero effective feedback delay was required for the Hopf instability and resulting oscillatory mode. Direct measurement of this delay, denoted *η* in the dimensional formulation and *τ* after nondimensionalization (see Supplementary Text S1), is difficult because it summarizes the finite response times of multiple steps involving signaling, cytoskeletal reorganization, force generation, and adhesion remodeling. Dimensionalizing and inverting the near-linear power law *T* = 6.88*τ* ^0.854^ obtained in the oscillatory regime gives *η* = *θ*_*R*_[*T*_dim_*/*(6.88*θ*_*R*_)]^1*/*0.854^, where *T*_dim_ is the dimensional oscillation period and *θ*_*R*_ is the common timescale governing the response and relaxation of both model variables. Reported timescales span tens of seconds to tens of minutes, depending on the process, measured quantity, and spatial scale [51]. Force-dependent FA growth [9, 52], integrin-mediated RhoA activation [10, 53], and RhoA-driven traction changes [14, 37] all occur within minutes. At individual FAs, FA assembly can precede FAK activation by approximately 43 s [18], and traction can precede FAK activation by 30–60 s [19], whereas adhesion-dependent RhoA regulation and mature FA persistence extend over tens of minutes [36, 54]. These observations suggest 1–100 min as a broad plausible range for *θ*_*R*_, although it was not measured directly. Using the approximately 10^2^ min recurrence timescale observed in representative local time courses (Fig. 1 and Fig. S2) as an illustrative proxy for *T*_dim_ then gives *η* ≈10–23 min. Independent estimates of *θ*_*R*_, combined with perturbations expected to alter feedback timing, could test whether the characteristic timescale changes as predicted by a delay-driven mechanism. More broadly, because feedback delays can alter stability and generate oscillatory or more complex dynamics, estimating the effective delay may help explain dynamical transitions in other feedback systems [21].

The bistable regime predicted by the model introduces history dependence: at the same parameter values, either the low- or high-activity state can persist depending on the system’s prior state. Such hysteresis permits state maintenance under modest changes while allowing switching after sufficiently large changes, resembling the balance between robustness and plasticity in biological regulation [55, 56]. A related mechanochemical model also yielded bistable contractility and traction hysteresis in response to time-dependent changes in substrate stiffness [25]. This prediction could be tested experimentally by slowly increasing and decreasing a factor expected to influence *α* and determining whether FA assembly and traction switch at direction-dependent thresholds. Possible approaches include real-time stiffness control [57], reversibly tunable hydrogels [58, 59], reversible control of integrin-binding ligand presentation [60], or gradual increases and decreases in the concentration of a reversible myosin-II inhibitor [43]. Because *A* and *F* are coarse-grained variables, these tests should focus on cell-level measures or spatial distributions of adhesion assembly and traction. If observed experimentally, such direction-dependent switching thresholds would support history-dependent mechanochemical switching in cell–ECM adhesion.

Numerous mathematical models have been developed to describe cell–ECM adhesion dynamics. Mechanochemical models have coupled adhesion formation to the contractility of stress fibers [22, 23], identified regimes of stress fiber– adhesion reorganization or collapse [61], and described force-dependent adhesion growth and maturation [62, 63]. Related signaling models have described Rac- and myosin-dependent regulation of protrusion and adhesion maturation [64], bidirectional signaling–mechanical coupling underlying intermittent leading-edge motility [24], and Rac/Rho-dependent vascular smooth muscle cell phenotypic plasticity [65]. Bistability or hysteresis has been predicted for receptor–ligand adhesion clusters [66], Rho-dependent cellular contractility [25], and mechanically coupled adhesion states arising from nonlinear linkage elasticity [67]. Motor–clutch loading can generate oscillatory traction [26], actin-network feedback has been linked to traction oscillations at FAs [27], and myosin-dependent motor–clutch dynamics can undergo a Hopf bifurcation to sustained force oscillations [68]. A recent preprint modeled delayed ECM displacement as negative feedback regulating nascent-adhesion turnover [69]. These studies provide mechanistic explanations for specific aspects of adhesion dynamics, including maturation, state switching, oscillation, and turnover. Within this broader modeling landscape, our minimal mechanochemical model interprets stable adhesion maintenance and dynamic remodeling in terms of three effective control parameters: feedback strength, feedback balance, and delay. The delay–period relation and direction-dependent switching thresholds provide experimentally testable predictions of the model.

Several limitations should be considered when interpreting these findings. The correspondence between the experimental temporal patterns and modeled dynamics remains qualitative. The proposed roles of RhoA-and Rac1-related signaling were not directly tested by measuring pathway activity or applying pathway-specific perturbations. In the model, pathway-specific kinetics, thresholds, and delays were represented by shared effective parameters. Extending the current nonspatial model by treating *β* as a dynamic field of local RhoA/Rac1 feedback balance could reveal whether spontaneous heterogeneity generates adhesion–force domains or mechanochemical waves that shape cell-wide force transmission. Experimentally, simultaneous measurements of adhesion, traction, and feedback signaling under controlled perturbations could test how feedback balance and timing shape adhesion dynamics across cell types and extracellular environments.

## IV. MATERIALS AND METHODS

### A. Cell culture

A7r5 rat aortic smooth muscle cells (CRL-1444, ATCC) were transfected with the pTagRFP–vinculin vector (FP372, Evrogen) to establish a stable RFP–vinculin-expressing cell line. Cells were maintained in Dulbecco’s modified Eagle medium containing 1.0 g*/*L glucose (Wako) supplemented with 10% heat-inactivated fetal bovine serum (SAFC Biosciences) and 1% penicillin–streptomycin (Wako) at 37 °C in a humidified atmosphere containing 5% CO_2_.

### B. Preparation of PDMS substrates

Polydimethylsiloxane (PDMS) silicone elastomer (SYLGARD 184, Dow Corning) was prepared at a base-to-curing-agent mass ratio of 50:1. After mixing, the PDMS mixture was degassed for 1 h, and 0.20 g was deposited onto each 35-mm glass-bottom dish with a 27-mm-diameter glass region (Iwaki). The PDMS was spin-coated, degassed, and then cured at 60 °C for 20 h. Micropatterns were generated as described previously [70] to maintain cells within the field of view during time-lapse imaging. Specifically, copper electron microscopy (EM) meshes (G201 and G203, EM Japan) were placed on the PDMS surface as shielding masks, and the uncovered square regions were exposed to oxygen plasma for 3 min at a chamber pressure below 20 Pa and a discharge current of 4 mA using a plasma generator (SEDE-P, Meiwafosis). The meshes were removed after plasma exposure, leaving square micropatterned regions of 120 × 120 *µ*m (G201) or 55 × 55 *µ*m (G203). Prior to cell seeding, the substrates were treated with 0.2% Pluronic F-127 (Invitrogen) for 1 h at 4 °C and subsequently with 0.1% gelatin (Sigma) for 3 h at 4 °C. Cells were then seeded onto the micropatterned PDMS substrates and allowed to spread overnight before time-lapse imaging.

### C. Live-cell imaging

More than 10 independent experiments were performed, and four representative recordings (R1–R4) were selected for the analyses presented here. The 120-*µ*m micropatterns were used for R1, R3, and R4, whereas the 55-*µ*m micropatterns were used for R2. Live-cell imaging was performed using an inverted microscope (IX-71, Olympus) equipped with a stage-top incubator (Tokai Hit) maintained at 37 °C and 5% CO_2_ and a camera (ORCA-R2, Hamamatsu), with image acquisition controlled using MetaMorph software (Molecular Devices). Using a 60 × oil-immersion objective, paired phase-contrast and RFP–vinculin fluorescence images were acquired sequentially at each time point with exposure times of 100 ms and 1000 ms, respectively. Images were acquired at 1-min intervals for R1 and 2-min intervals for R2–R4.

### D. Wrinkle force microscopy

We previously developed SW-UNet, a convolutional neural network that extracts cell-generated wrinkles in plasma-treated PDMS substrates from phase-contrast images [71]. Using these extracted wrinkle images, we subsequently developed wrinkle force microscopy (WFM) to reconstruct cellular traction-stress fields [28]. The WFM model was trained on paired data comprising SW-UNet-extracted wrinkle images of A7r5 cells and corresponding traction-stress fields measured by conventional traction force microscopy (TFM). Cell-generated substrate wrinkles in the present phase-contrast recordings were extracted at each time point using SW-UNet, and each extracted wrinkle image was provided as input to the pretrained WFM model. The model returned a 26×26 grid of two-dimensional traction-stressvectors (*T*_*x*_, *T*_*y*_). At each grid point, the WFM-derived traction-stress magnitude was calculated as 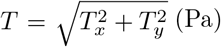.

### E. Image and data analysis

Four time-lapse datasets, R1–R4, were analyzed using RFP–vinculin image sequences and traction-stress vector fields. At each time point, the traction analysis yielded a 26 × 26 grid of traction-stress vectors, and the corresponding RFP image crop was divided into matching regions (Fig. S1). At each grid position, the local RFP–vinculin signal was calculated as the mean pixel intensity within the corresponding grid cell (arbitrary units), and local traction stress as the vector magnitude. The two signals were paired by frame number and grid position, and all grid positions were included without spatial masking.

Each time series was smoothed over its complete duration using a centered arithmetic moving mean spanning ± 10 min. For the spatial amplitude analysis, overlapping 240-min windows were advanced in 30-min steps and retained only when the centered moving mean was defined at every time point and both signals were finite. The smoothed RFP–vinculin and traction signals were separately linearly detrended within each window using ordinary least-squares (OLS) regression. For each recording and signal, the recording-wide median *M*_*X*_ was calculated from the smoothed values pooled over all grid positions and all time points for which the centered moving mean was defined. For each grid position, the relative deviation within a window was expressed as a percentage of this recording-wide signal level and calculated as

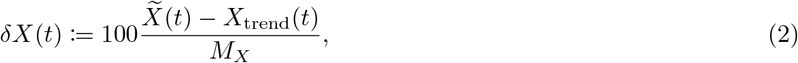

where 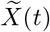 is the smoothed local RFP–vinculin or traction signal and *X*_trend_(*t*) is its fitted linear trend within the window. Joint trajectories were constructed by plotting the traction deviation against the RFP–vinculin deviation at corresponding time points in chronological order. For Fig. S2 and Fig. S3, the raw and smoothed local signals were detrended separately within each displayed window and normalized by the corresponding recording-wide median *M*_*X*_ . Smoothing sensitivity was evaluated using centered moving means with durations of 15, 20, and 30 min, using the *M*_*X*_ obtained from the centered ±10-min condition for all three conditions (Fig. S3).

### F. Quantification and spatial mapping of fluctuation amplitudes

For each 240-min window, the fluctuation amplitude of each signal was defined as the 10th-to-90th percentile range of its relative deviation,

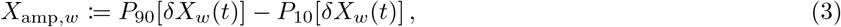

where *X* represents RFP–vinculin intensity or traction stress, and *w* denotes the analysis window. The amplitude was expressed as a percentage of the corresponding recording-wide median signal level. For each grid position, the median amplitude across all included windows was used as its representative fluctuation amplitude. The spatial maps in Fig. 2A and B display the grid-level median amplitudes at their corresponding sampling positions without additional spatial smoothing or interpolation.

Gaussian kernel-density estimates of the grid-level median amplitudes were calculated separately for R1–R4 in log_10_ amplitude space. A common reference bandwidth was determined for each signal using the robust rule-of-thumb estimator

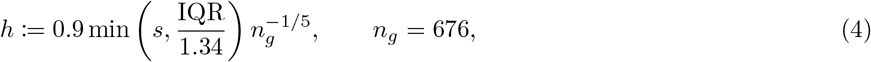

where *s* and IQR are the standard deviation and interquartile range of the pooled transformed amplitudes and *n*_*g*_ is the number of grid positions in each recording. The aggregate density labeled All was calculated as the arithmetic mean of the four recording-specific densities so that R1–R4 contributed equally (Fig. 2C and D). Normalization sensitivity was examined by replacing the recording-wide median *M*_*X*_ with the median of the corresponding smoothed local signal within each window while retaining the same residuals and amplitude definition. The resulting grid-level estimates were compared within each recording using Spearman’s rank correlation coefficient, and the same density-estimation procedure was used to compare their distributions (Fig. S4). Distribution sensitivity was evaluated by comparing bandwidths of 0.75*h*, 1.00*h*, and 1.25*h*, and by comparing the standard median across 240-min windows with the median across 360-min windows advanced in the same 30-min steps and with the mean, 25th percentile, and 75th percentile across 240-min windows (Fig. S5).

### G. Mechanochemical model formulation

We formulated a dimensionless mechanochemical model of reciprocal feedback between focal adhesion assembly and actomyosin tensile force to examine how this feedback architecture can generate stable and oscillatory modes of cell–ECM adhesion dynamics (Fig. 3A). The resulting delay differential equations (DDEs) are given in Eq. (1). The dimensionless state variables *A*(*t*) and *F* (*t*) represent focal adhesion assembly and actomyosin tensile force, respectively. All three feedback interactions were represented by delayed Hill functions, with an increasing force-to-adhesion function for force-dependent FA assembly and maturation and increasing RhoA-related and decreasing Rac1-related functions for the opposing adhesion-to-force feedbacks. The linear loss terms represent effective FA turnover and relaxation of AM tensile force. Force-dependent FA maturation is supported by mechanical measurements, force-application experiments, and molecular studies of mechanosensitive adhesion reinforcement [3, 9, 12, 34, 42, 43, 72]. Adhesion-associated signaling regulates both RhoA and Rac1 [10, 33, 73, 74]. Experimental and modeling studies further identify molecular routes for reciprocal inhibition and coordinated RhoA–Rac1 dynamics [16, 75–77]. RhoA- and Rac1-related signaling can in turn regulate AM contractility [8, 14, 78–80].

The mechanochemical coupling strength *α* sets the overall strength of the reciprocal feedback between FA assembly and AM tensile force. The feedback-strength ratio *β* gives the strength of the Rac1-related negative feedback relative to the RhoA-related positive feedback. The parameter *τ* denotes the effective feedback delay shared by the three interactions and represents the accumulated latency of signaling, cytoskeletal reorganization, force generation, and adhesion remodeling [20, 21, 81].

For *τ* = 0, the model reduces to an ordinary differential equation (ODE) system, whereas for *τ >* 0, it requires history functions defined on *t* ∈ [− *τ*, 0], which were taken to be positive [82]. The model was used to examine how the feedback architecture organizes dynamical modes without fitting its parameters to the experimental RFP–vinculin or traction time series. The biological rationale, dimensional formulation, nondimensionalization, and reduction to the three-parameter model are described in detail in Supplementary Text S1.

### H. Linear stability and bifurcation analysis

Because the delayed and current states coincide at a fixed point, positive fixed points (*A*^∗^, *F* ^∗^) were determined numerically for each (*α, β*) combination from the *τ* -independent steady-state equations obtained by setting d*A/*d*t* = d*F/*d*t* = 0 in Eq. (1). We linearized the model around each fixed point and defined

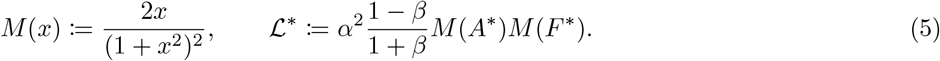

Substituting perturbations of the form (*δA*(*t*), *δF* (*t*)) = (*u, v*)*e*^*λt*^ into the linearized system gave the characteristic equation

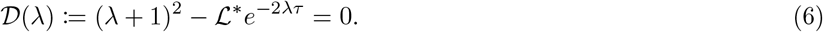

A fixed point was classified as linearly asymptotically stable when every characteristic root satisfied Re(*λ*) *<* 0.

The saddle-node boundary was identified by setting *λ* = 0 in the characteristic equation, giving ℒ^∗^ = 1 and requiring 0 *< β <* 1. Because both the fixed-point equations and the zero-root condition are independent of *τ*, the saddle-node boundary is also independent of the feedback delay. The boundary was constructed by mapping positive fixed-point coordinates satisfying ℒ^∗^ = 1 to the (*α, β*) plane using the fixed-point parameterization. On this boundary, the zero root was simple because *∂ D/∂λ* = 2(1 + *τ* ) *>* 0 at *λ* = 0 and ℒ^∗^ = 1. Fixed-point branch and nullcline analyses confirmed that these zero-root points were saddle-node folds.

The linear Hopf boundary was identified by substituting *λ* = i*ω*, with *ω >* 0, into Eq. (6). Formal purely imaginary solutions for *β <* 1 require ℒ^∗^ *>* 1, where a positive real characteristic root already makes the fixed point unstable. No Hopf solution exists at *β* = 1. The first loss of fixed-point stability for *β >* 1 was therefore determined by

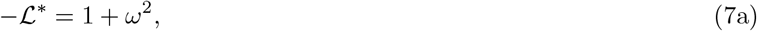

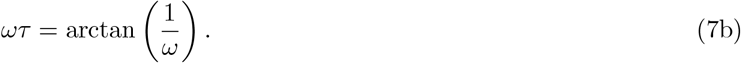

For each prescribed *τ >* 0, *ω* was first determined from Eq. (7b), and positive fixed-point coordinates satisfying Eq. (7a) were then mapped to the (*α, β*) plane. The phase condition selects the smallest positive critical delay and hence the first loss of fixed-point stability as *τ* increases. No Hopf bifurcation occurs at *τ* = 0. The purely imaginary roots were simple, and d Re(*λ*)*/*d*τ >* 0 on the boundary, confirming that they crossed from the stable side to the unstable side as *τ* increased. Direct numerical integration was used to verify convergence to sustained periodic solutions at the examined parameter conditions on the oscillatory side of the boundary. The complete fixed-point parameterization and derivations of the characteristic equation and the saddle-node and Hopf bifurcation conditions are provided in Supplementary Text S2.

### I. Numerical simulations and trajectory analysis

#### Numerical integration and history functions

For *τ >* 0, the DDEs were integrated using dde23 in MATLAB R2024b (The MathWorks, Inc.). The relative tolerance was set to 10^−6^, the absolute tolerance used the MATLAB default of 10^−6^, and the maximum step size was set to 0.05. Unless otherwise stated, the simulations used the constant history function

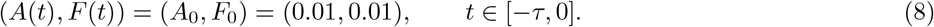

The bistable trajectory shown as case iv in Fig. 4B was additionally simulated using the constant history (*A*_0_, *F*_0_) = (0.3, 0.3). For the *τ* = 0 trajectory shown as case i in Fig. 4B, the model was integrated as an ODE system from (*A*(0), *F* (0)) = (0.01, 0.01) using the DOP853 method implemented in SciPy’s solve ivp, with relative and absolute tolerances of 10^−10^ and 10^−12^, respectively.

#### Construction of dynamical-regime maps

The phase diagrams in Fig. 3B and Fig. 4A were constructed using the saddle-node and Hopf bifurcation curves identified by the linear stability analysis. The delay-independent saddle-node curve was used for all values of *τ*, whereas the first Hopf boundary was calculated separately for *τ* = 0.5, 1, and 5. For each finite delay, the lower-*β* envelope of the Hopf solutions was retained as the boundary corresponding to the first loss of fixed-point stability. The two saddle-node branches bounded the bistable region, and the first Hopf boundary delimited the oscillatory region. Because *A*^∗^ and *F* ^∗^ varied continuously across the remaining monostable parameter space without additional bifurcations, descriptive subdivisions were defined operationally using the near-vertical saddle-node branch and the *β* coordinate of its minimum-*α* endpoint. These subdivisions were labeled as the inactive, low-activity stable, high-activity stable, and adhesion-dominant regions according to the qualitative magnitudes and relative balance of *A*^∗^ and *F* ^∗^ (Fig. S8).

#### Fixed-point and phase-plane analysis

For the nullcline and phase-plane analysis in Fig. 4C, the nondelayed system (*τ* = 0) was evaluated at *β* = 0.01 and *α* = 1, 3, and 15. The *A* and *F* nullclines were obtained by setting d*A/*d*t* = 0 and d*F/*d*t* = 0, respectively, and the positive fixed points were identified numerically as their intersections using SciPy’s fsolve from multiple initial guesses. The linear stability of each fixed point was determined from the eigenvalues of the corresponding 2 × 2 Jacobian. Streamlines were calculated from the ODE vector field and displayed in (log_10_ *A*, log_10_ *F* ) coordinates.

For the steady-state branch analysis in Fig. 4D, the fixed-point equations were solved at 1,000 linearly spaced values of *α* over 0.05 ≤ *α* ≤ 20 with *β* = 0.01. The resulting solutions were separated into lower stable, upper stable, and unstable branches. At *β* = 0.01, these stability assignments are independent of *τ* under the characteristic-root conditions derived above. The saddle-node points were identified using the zero-root condition ℒ^∗^ = 1. Hysteresis under quasistatic variation of *α* was inferred from the coexistence of the two stable branches and their distinct saddle-node switching thresholds without simulating a finite-rate parameter sweep. For Fig. S8, fixed points across the sampled (*α, β*) grid were grouped into the corresponding monostable and bistable solution families.

#### Parameter-grid simulations and trajectory preprocessing

Oscillatory properties were analyzed over a parameter grid comprising *α* ∈ {0, 1, …, 20}, *β*_*j*_ = 10^2+*j/*3^ for *j* ∈ {0, 1, …, 9 }, and *τ*_*k*_ = 10^−1+*k/*13^ for *k* ∈ {0, 1, …, 39 }, giving 8,400 parameter combinations. The value *α* = 0 was included only as the zero-feedback boundary case; the biologically relevant domain otherwise assumes *α >* 0. Each trajectory was integrated to *t* = 5000 using the constant history (*A*_0_, *F*_0_) = (0.01, 0.01). The interval *t >* 1500 was retained for post-transient analysis. Because dde23 returns solutions at adaptively selected time points, post-transient trajectories containing more than 30,000 points were reduced to 30,000 approximately evenly indexed samples. The retained *A* and *F* values were then linearly interpolated onto a common evenly spaced time grid before calculating the period, amplitudes, and phase difference.

#### Quantification of oscillatory properties

To identify trajectories with sufficient post-transient variation for numerical quantification of oscillatory properties, we required the standard deviation of the original post-transient *A*(*t*) samples to exceed 0.1 before downsampling and interpolation. Only trajectories satisfying this criterion and yielding finite estimates of the period, both amplitudes, and phase difference were included. The oscillation period *T* was defined as the lag of the first positive-lag peak in the autocorrelation of the mean-centered, resampled *A*(*t*) signal. The autocorrelation peak was required to reach at least 10% of the autocorrelation maximum. For *X*(*t*) ∈ {*A*(*t*), *F* (*t*)}, the oscillation amplitude was defined as

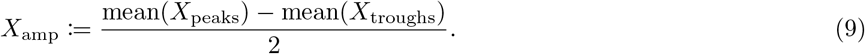

Local maxima were detected using SciPy’s find peaks routine, and troughs were detected by applying the same routine to −*X*(*t*). At least two peaks and two troughs were required for amplitude calculation.

The time difference Δ*t* was defined as the lag maximizing the unnormalized full cross-correlation of the mean-centered *F* (*t*) signal with the mean-centered *A*(*t*) signal, calculated in that order. The circular phase difference was defined as

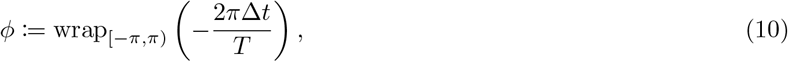

where wrap_[−*π,π*)_ maps the angle to the principal interval [−*π, π*). With this convention, *ϕ >* 0 indicates that *F* leads For a set of *n* phase differences *ϕ*_*j*_, the mean resultant vector was calculated as 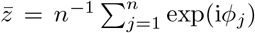. The circular mean 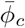, mean resultant length *R*, and circular standard deviation *σ*_*c*_ were then calculated as

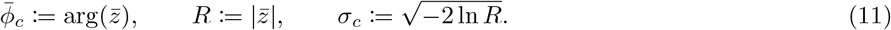

Here, arg 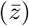 and 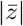 denote the angle and magnitude of the mean resultant vector, respectively. For the parameter-resolved phase summaries, 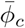 and *σ*_*c*_ were calculated separately for each fixed *α* and each fixed *β*.

The pooled delay–period relationship was estimated by unweighted OLS regression of log_10_ *T* against log_10_ *τ* . Separate delay–period fits were calculated for each fixed *α* and each fixed *β*. The 95% *t*-based intervals for the fitted exponents were calculated from their OLS standard errors. For each fixed (*β, τ* ) combination, *A*_amp_ and *F*_amp_ were regressed separately against *α* using at least three distinct *α* values that satisfied the trajectory-selection criteria. The fitted slopes quantified amplitude sensitivity to *α*, and *R*^2^ quantified the goodness of the corresponding linear fit. The amplitude ratio was calculated as *A*_amp_*/F*_amp_.

For the representative trajectories in Fig. 5D, the final complete post-transient cycle was delimited by the last two peaks of the original post-transient *A*(*t*) signal. The displayed normalized cycle time, denoted *t/T*, was calculated as (*t* − *t*_start_)*/*(*t*_end_ − *t*_start_).

## DATA AVAILABILITY

The processed experimental and numerical source data underlying the quantitative results in the main and Supplementary figures are provided as Source Data with this manuscript.

## CODE AVAILABILITY

The custom MATLAB and Python code used for numerical simulations, bifurcation analysis, source-data validation, and figure generation is provided as Supplementary Software with this manuscript and will be made publicly available at https://github.com/EijiMatsumoto/cell-matrix-adhesion-delay-model upon acceptance.

## ACKNOWLEDGEMENTS

We thank Honghan Li and Daiki Matsunaga for providing the WFM implementation in a form suitable for the present analysis.

## AUTHOR CONTRIBUTIONS

E.M. and S.D. conceptualized the study. T.A., T.S.M., and S.Y. performed the experiments and acquired the time-lapse data. E.M. developed the mathematical model, analyzed the experimental data, performed the theoretical and computational analyses, prepared the figures, and led the writing of the manuscript. S.D. conceived and supervised the experimental data acquisition, provided resources, supervised the project, and contributed to the interpretation and writing of the manuscript. All authors reviewed and edited the manuscript. E.M. and S.D. acquired funding.

## USE OF GENERATIVE AI

During preparation of this manuscript, the authors used ChatGPT and Codex (OpenAI) to assist with language editing and text revision, preliminary literature exploration, and the development and debugging of analysis and simulation code. The authors reviewed and revised all AI-assisted outputs and tested the resulting code. All scientific content, analyses, interpretations, and cited references were independently verified by the authors, who take full responsibility for the manuscript.

## SUPPLEMENTARY INFORMATION

### Supplementary Figures

**FIG. S1.**
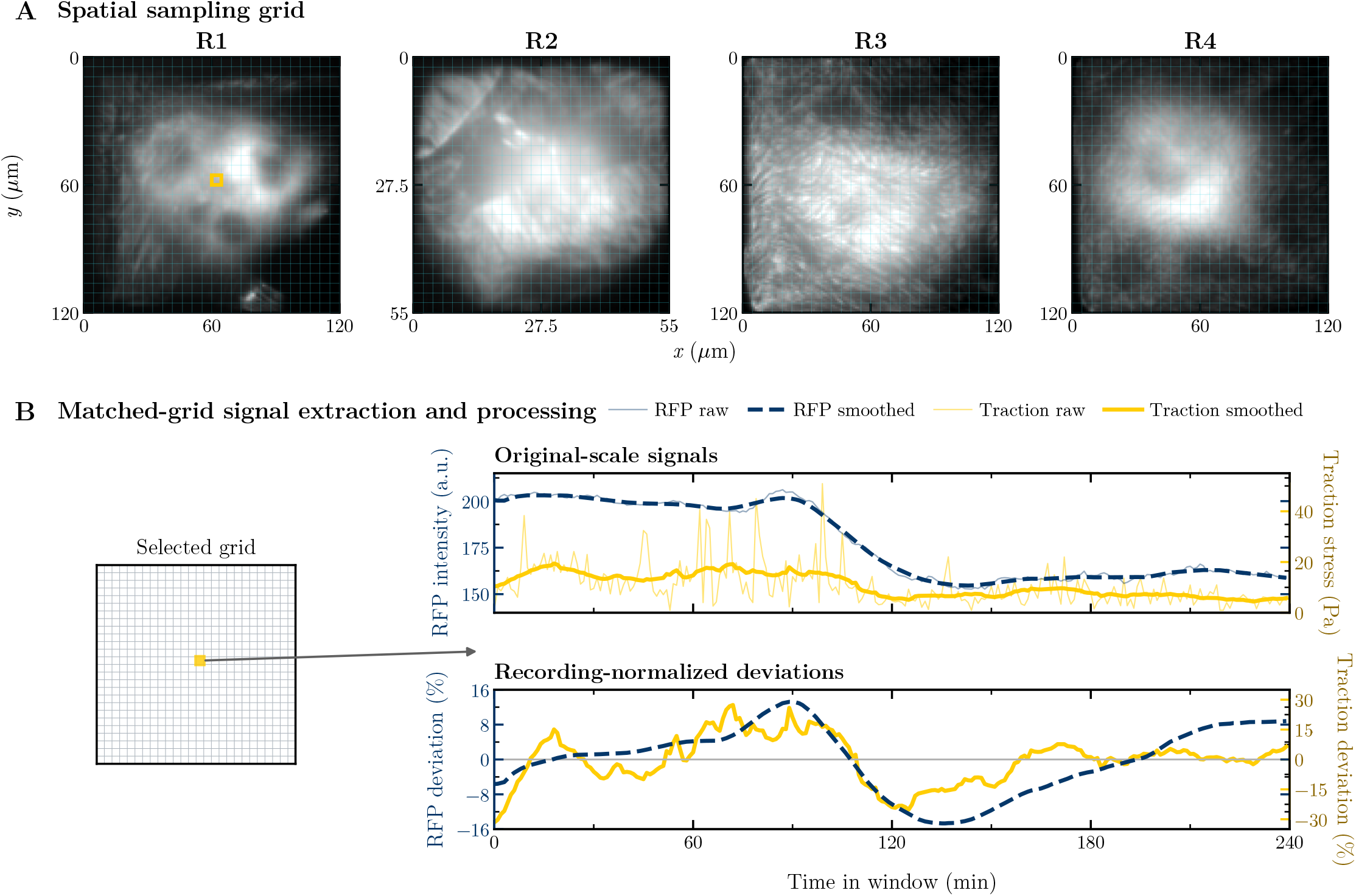
Matched spatial sampling pairs local adhesion and traction signals for temporal analysis. Within each of four time-lapse datasets, corresponding RFP–vinculin and traction signals were extracted from a matched 26 × 26 grid and processed into recording-normalized deviations over 240-min windows. **(A)** RFP–vinculin images averaged over each complete time series are shown with the corresponding spatial sampling grids for R1–R4. The highlighted location in R1 identifies the grid position illustrated in panel B. **(B)** RFP–vinculin intensity and traction stress from the matched grid position are shown over an illustrative 240-min window. The upper traces show the original-scale raw signals and centered ± 10-min moving averages calculated over the complete time series before window extraction. The lower traces show recording-normalized deviations calculated independently for each channel as 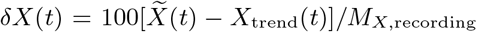_,recording_, where 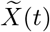 is the smoothed signal, *X*_trend_(*t*) is its fitted linear trend within the window, and *M*_*X*,recording_ is the corresponding recording-wide median signal level.

**FIG. S2.**
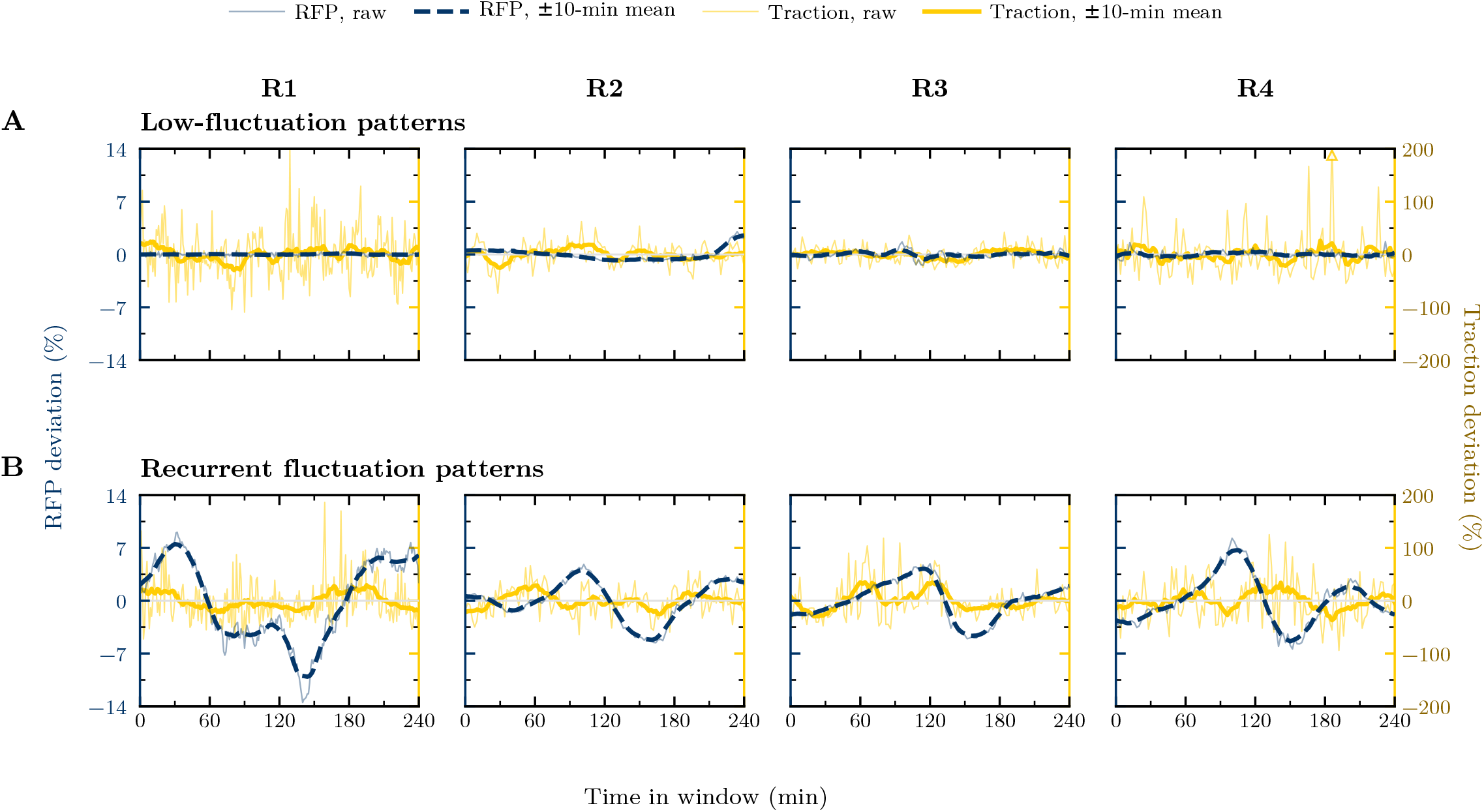
Representative examples show low-fluctuation and recurrent fluctuation patterns across time-lapse datasets. One low-fluctuation pattern and one recurrent fluctuation pattern from each time-lapse dataset are shown over 240-min windows using common axes and the same signal-processing procedure as in Fig. 1. **(A)** Low-fluctuation patterns from R1–R4 show comparatively limited deviations in the smoothed RFP–vinculin and traction signals. **(B)** Recurrent fluctuation patterns from R1–R4 show increases and decreases in RFP–vinculin over the displayed windows, with corresponding traction signals that also vary over time. The R2 example in panel A and the R1 example in panel B correspond to the low-fluctuation and recurrent fluctuation patterns shown in Fig. 1, respectively. Thin and thick lines show deviations calculated from the raw and smoothed signals, respectively, and an open triangle marks the raw traction deviation outside the displayed ±200% range.

**FIG. S3.**
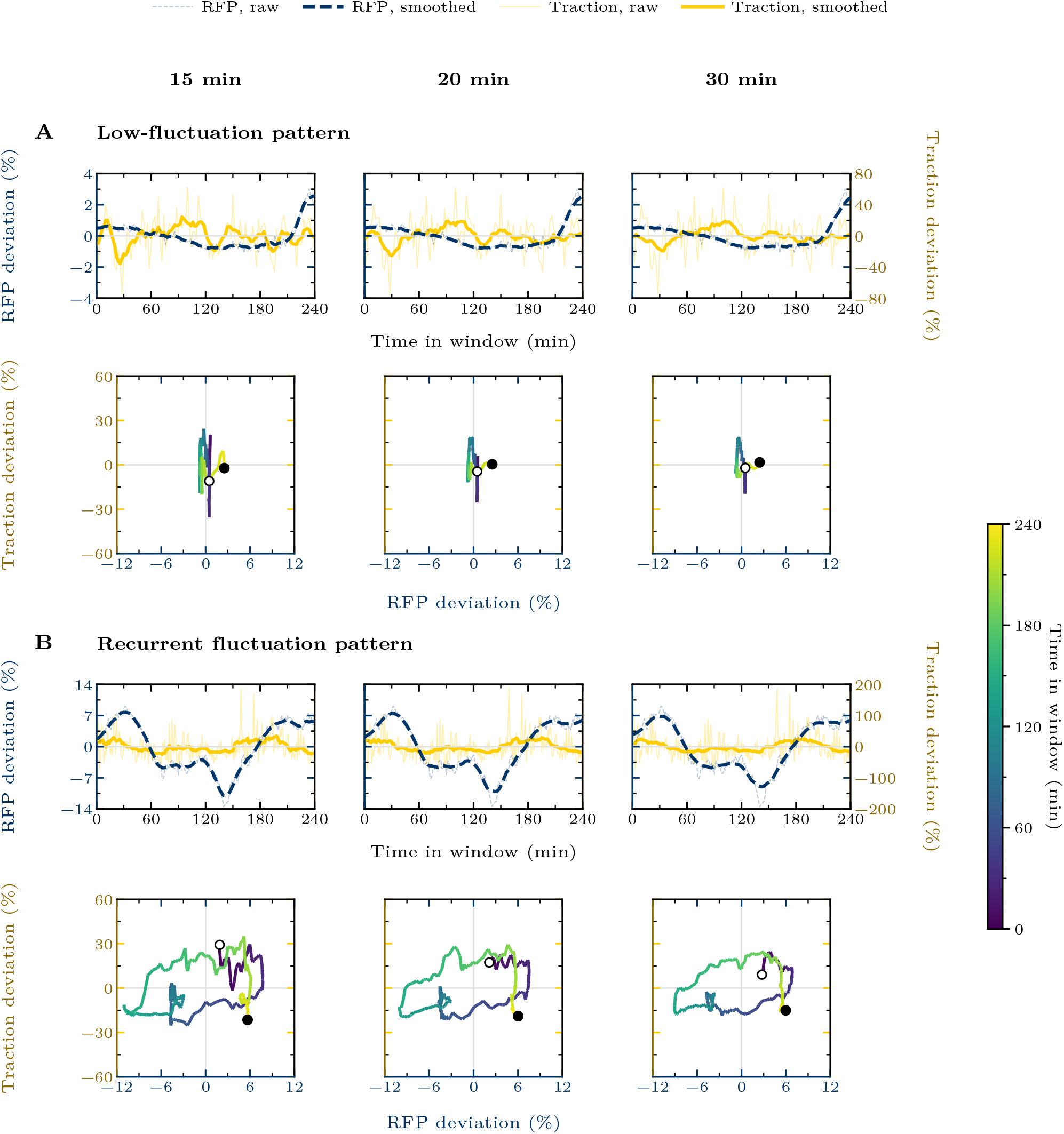
Representative temporal patterns are qualitatively preserved across smoothing windows. The low-fluctuation and recurrent fluctuation patterns in Fig. 1 retain similar waveform structures and joint-trajectory organization across the examined smoothing conditions, although stronger smoothing reduces fluctuation amplitudes, particularly for traction. **(A and B)** Across the three smoothing conditions, the low-fluctuation pattern retains limited RFP–vinculin and traction deviations and compact joint trajectories (A), whereas the recurrent fluctuation pattern continues to show increases and decreases in both signals and expanded joint trajectories (B). Columns indicate the nominal centered smoothing windows, and the 20-min condition corresponds to the centered ±10-min moving average used in Fig. 1.

**FIG. S4.**
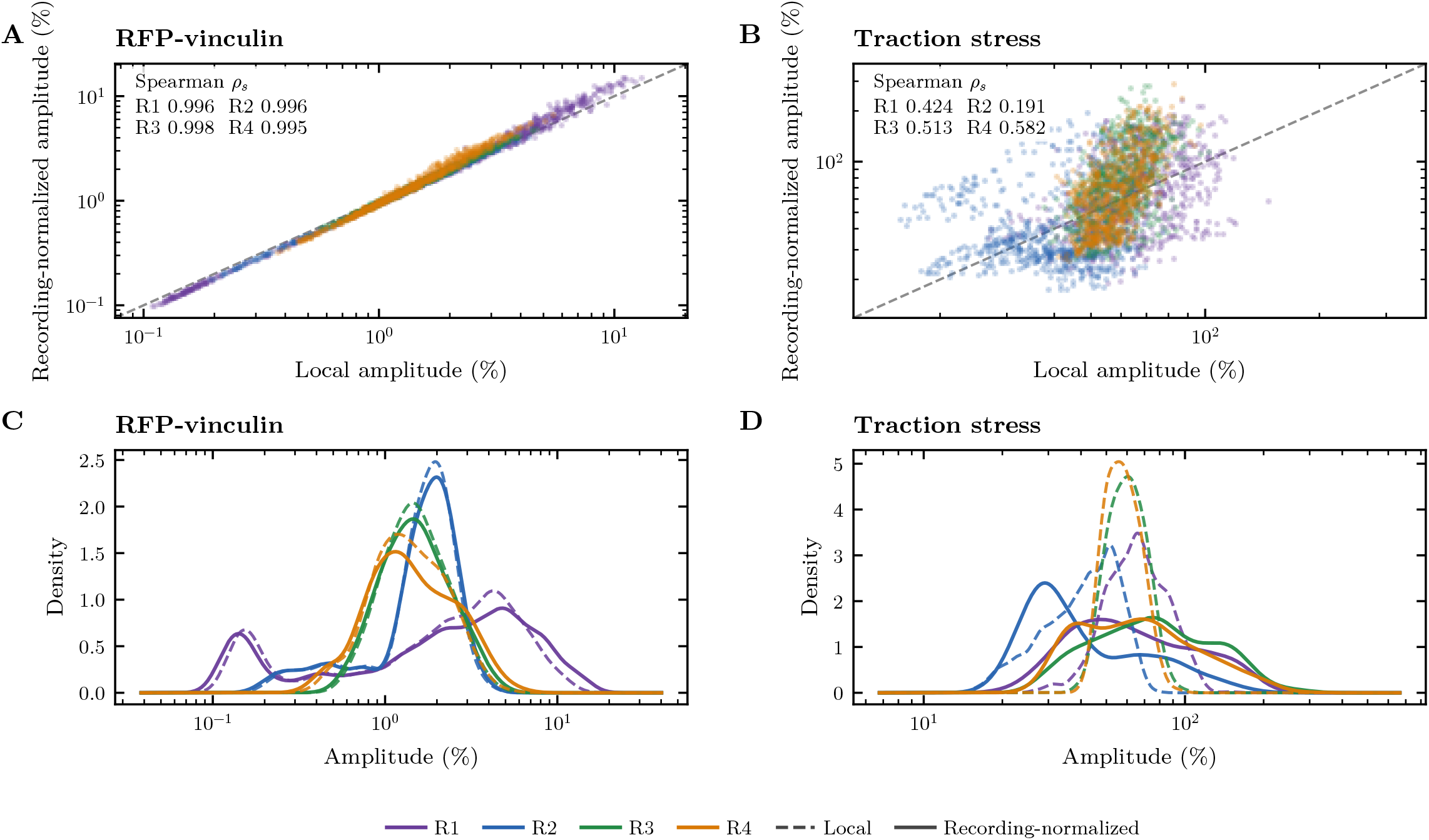
Broad RFP–vinculin and traction-stress amplitude variation persists across normalization choices. Window-local and dataset-wide median normalization yield positively correlated amplitude estimates in both channels, with substantially stronger agreement for RFP–vinculin than for traction stress. **(A and B)** Recording-normalized amplitudes are plotted against locally normalized amplitudes for RFP–vinculin (A) and traction stress (B), with each point representing one grid position and colors identifying R1–R4. Both normalization schemes use the same 10th-to-90th percentile range of the linearly detrended signal as the numerator, with local amplitudes divided by the corresponding window median and recording-normalized amplitudes by the dataset-wide median signal level. At each grid position, the median amplitude across the analyzed 240-min windows is shown. The dashed line denotes equality, and the annotations report the dataset-specific Spearman rank correlation coefficients *ρ*_*s*_. **(C and D)** Kernel-density estimates compare the corresponding local and recording-normalized amplitude distributions for RFP–vinculin (C) and traction stress (D). Dashed and solid curves denote local and recording-normalized amplitudes, respectively.

**FIG. S5.**
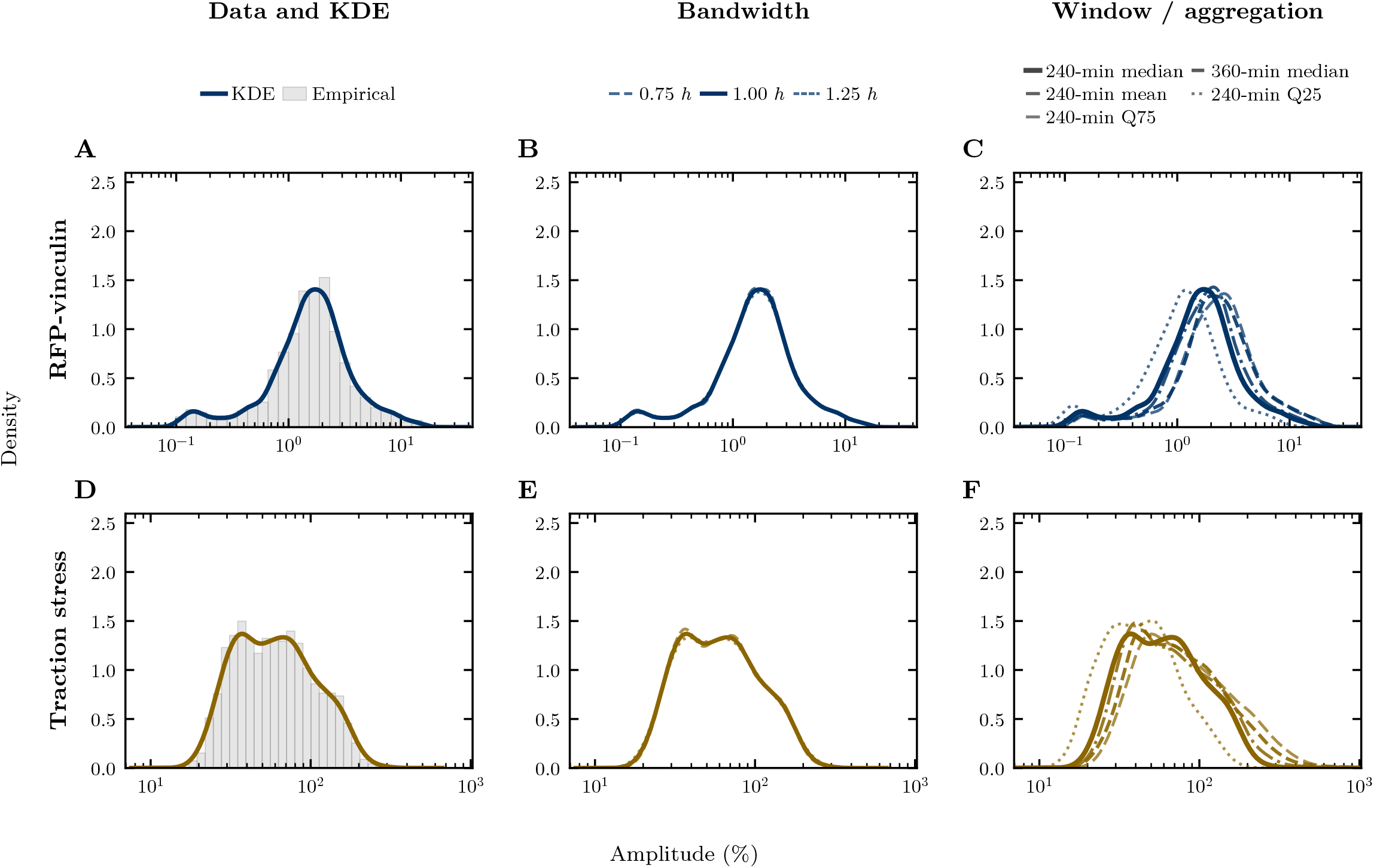
Broad RFP–vinculin and traction-stress amplitude distributions persist across analysis choices. Both signals retain broad amplitude distributions when kernel-density bandwidth, window duration, and within-grid aggregation are varied. **(A and D)** Empirical distributions and their kernel density estimates (KDEs) are shown for the RFP–vinculin (A) and traction-stress (D) amplitudes used in Fig. 2. At each grid position, amplitude was summarized by the median across 240-min windows, and R1–R4 were combined with equal dataset weights. **(B and E)** KDEs are compared using bandwidths of 0.75 *h*, 1.00 *h*, and 1.25 *h*, where *h* denotes the bandwidth used in Fig. 2. **(C and F)** The 240-min median distribution used in Fig. 2 is compared with the 360-min median and with the mean, 25th-percentile, and 75th-percentile summaries across 240-min windows. Kernel-density estimation was performed on log_10_-transformed amplitudes. The position and prominence of secondary maxima vary among these conditions.

**FIG. S6.**
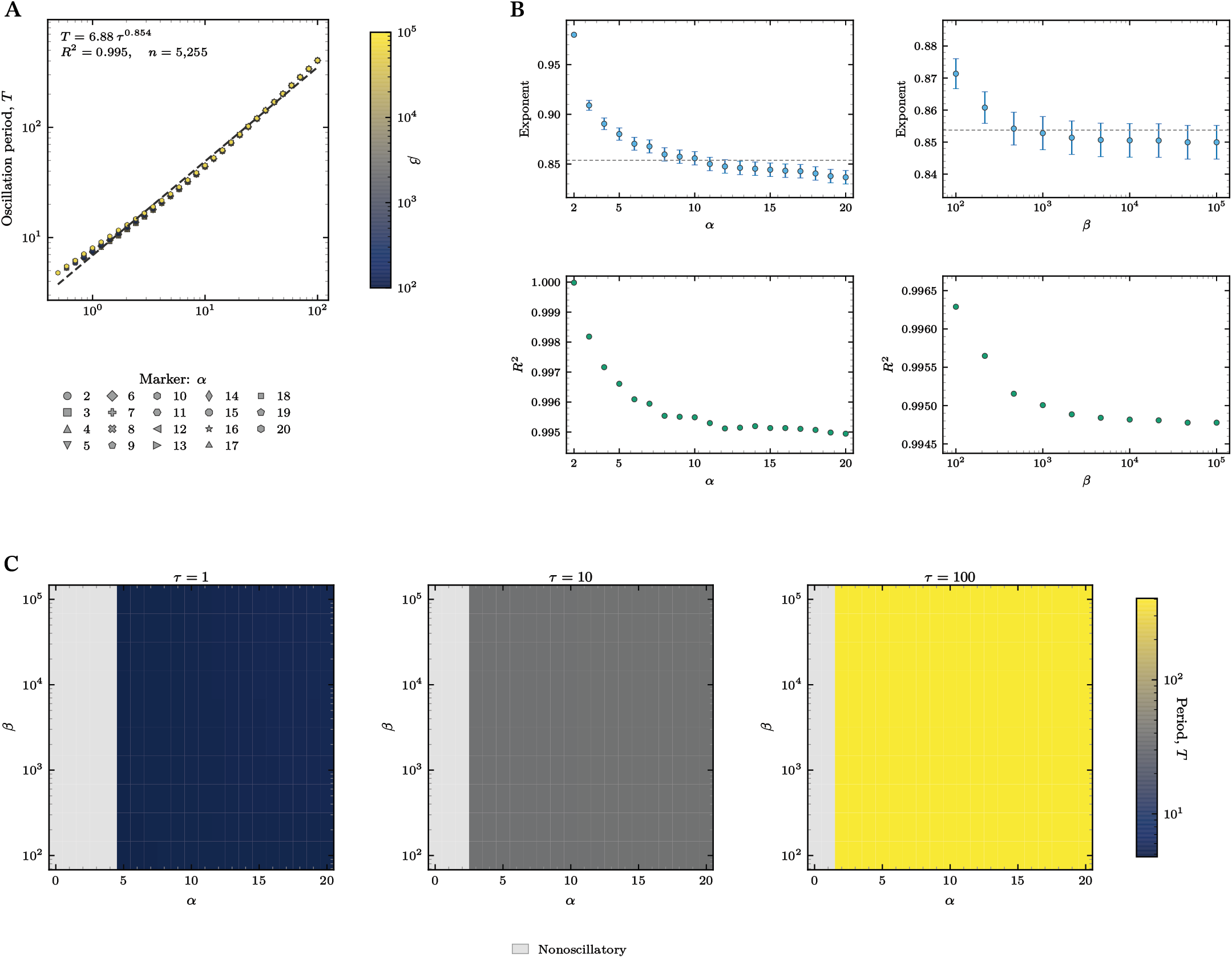
Delay–period scaling remains consistent across feedback parameters. Pooled and parameter-stratified fits, together with period maps at selected delays, show that *τ* primarily determines oscillation period while *α* and *β* introduce smaller systematic variations. **(A)** Oscillation period *T* is plotted against feedback delay *τ* for parameter combinations that produced sustained oscillations, with marker shape and color denoting *α* and *β*, respectively. The black dashed line shows the pooled power-law fit *T* = 6.88*τ* ^0.854^, with coefficient of determination *R*^2^ = 0.995. **(B)** Exponents and *R*^2^ values from separate power-law fits of *T* versus *τ* are shown for each fixed *α* (left) and each fixed *β* (right). Error bars show 95% *t* intervals derived from the ordinary least-squares standard errors of the fitted exponents, and the horizontal dashed line marks the pooled exponent from panel A. **(C)** Oscillation periods are mapped in the (*α, β*) plane at *τ* = 1, 10, and 100.

**FIG. S7.**
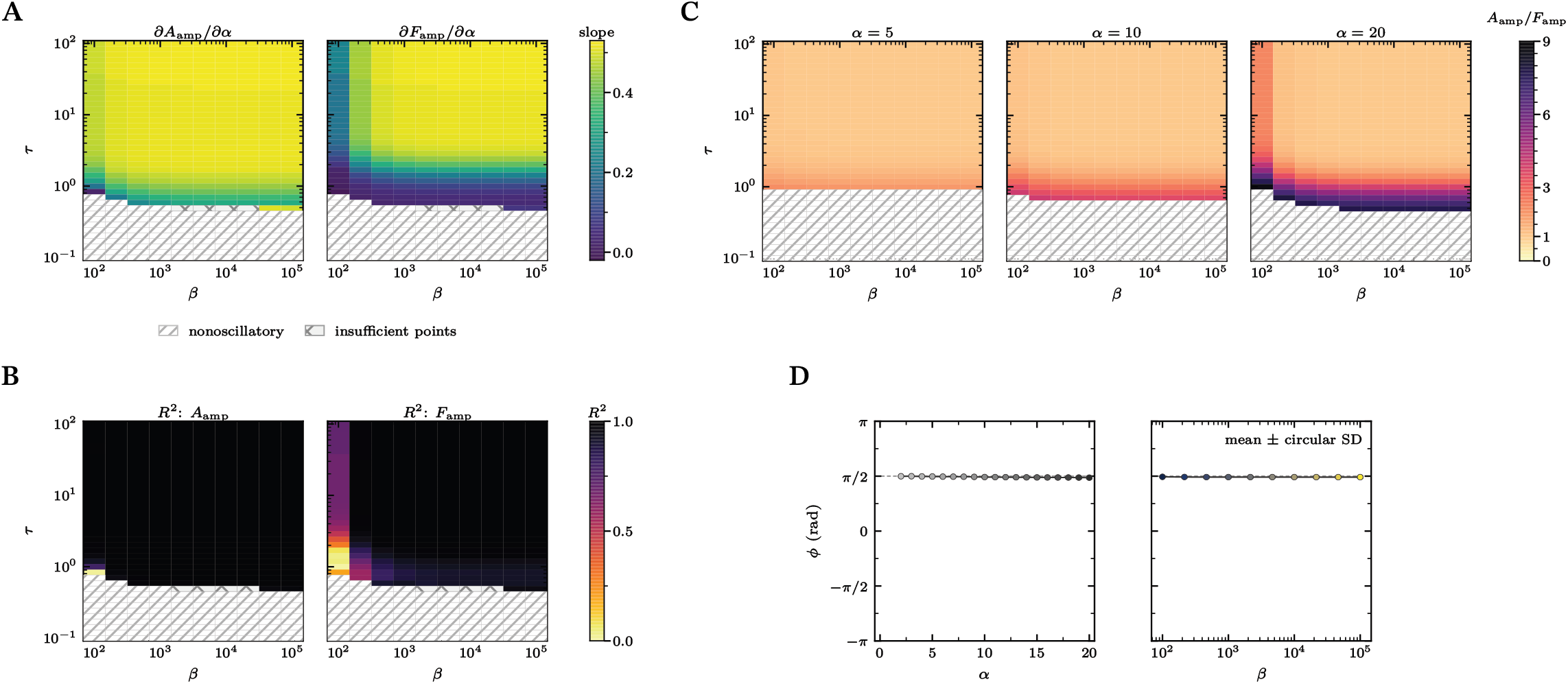
Feedback parameters and delay jointly shape oscillation amplitudes while the phase relationship remains constrained. Amplitude sensitivity to *α* and the relative amplitudes of *A* and *F* vary with *β* and *τ*, whereas the circular phase difference remains concentrated near *π/*2. **(A)** For each fixed (*β, τ* ) combination, the partial sensitivities *∂A*_amp_*/∂α* and *∂F*_amp_*/∂α* were estimated as the slopes of separate ordinary least-squares regressions against *α*. **(B)** For each regression in panel A, the coefficient of determination *R*^2^ quantifies the fraction of amplitude variation across *α* explained by the fitted linear relationship and therefore indicates how adequately the corresponding slope summarizes the *α* dependence. In panels A and B, diagonal hatching denotes nonoscillatory combinations, and cross-hatching denotes combinations with fewer than three oscillatory *α* values. **(C)** The amplitude ratio *A*_amp_*/F*_amp_ is mapped in the (*β, τ* ) plane at *α* = 5, 10, and 20, with hatched regions indicating nonoscillatory combinations. **(D)** Circular means of the phase difference *ϕ* are shown for each *α* (left) and each *β* (right), with error bars denoting circular standard deviations across the corresponding oscillatory parameter combinations. Positive *ϕ* indicates that *F* leads *A*, and the gray dashed line marks *π/*2.

**FIG. S8.**
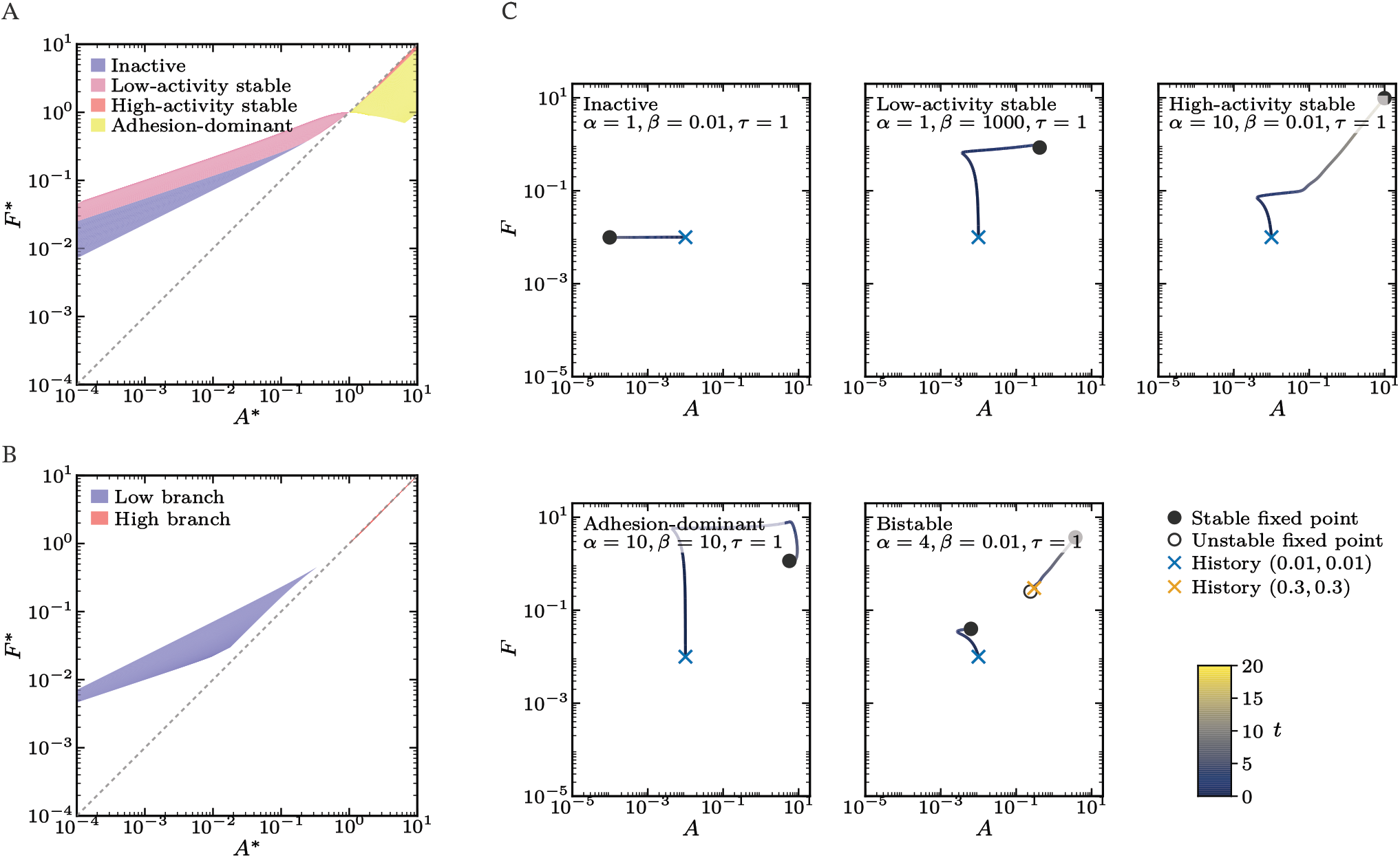
Fixed-point organization distinguishes nonoscillatory behaviors and permits history-dependent attractor selection. Fixed-point surfaces and representative trajectories show how the nonoscillatory fixed-point behaviors are organized in (*A, F* ) space and how different histories select alternative stable attractors in the bistable regime. **(A)** Colored surfaces show the stable fixed points of the four monostable subdivisions (inactive, low-activity stable, high-activity stable, and adhesion-dominant) across the sampled (*α, β*) parameter grid in raw (*A*^*∗*^, *F*^*∗*^) coordinates. **(B)** The low and high stable fixed-point branches of the bistable regime are shown in the same (*A*^*∗*^, *F*^*∗*^) coordinates. In panels A and B, the gray dashed line marks *F*^*∗*^ = *A*^*∗*^. **(C)** Representative trajectories at *τ* = 1 are shown for the indicated inactive, low-activity stable, high-activity stable, adhesion-dominant, and bistable conditions, with color denoting time over 0 ≤ *t* ≤ 20. Filled and open circles denote linearly stable and unstable fixed points, respectively, and colored × symbols mark the prescribed constant histories. The history (*A*_0_, *F*_0_) = (0.01, 0.01) is used for all five conditions, and the bistable condition additionally includes (0.3, 0.3); the two histories converge to different stable fixed points.

## SUPPLEMENTARY TEXT S1. BIOLOGICAL RATIONALE AND FORMULATION OF THE MECHANOCHEMICAL MODEL

### Biological rationale for the feedback architecture

The model represents reciprocal mechanochemical feedback between focal adhesion (FA) assembly and actomyosin (AM) tensile force using two effective state variables (Fig. 3A). The variable *A* denotes FA assembly, and the variable *F* denotes AM tensile force. Both variables represent collective states arising from multiple molecular and structural processes. In one direction of the feedback, AM-generated tension transmitted to FAs promotes their growth and reinforcement [1–3]. Cell-generated traction was closely associated with FA assembly [4], and experiments applying local mechanical force directly demonstrated FA enlargement [5]. At the molecular scale, tensile loading of talin exposes vinculin-binding sites, while force transmission through talin and vinculin strengthens coupling between integrins and the AM cytoskeleton [6–10]. Myosin II activity promotes FAK/Src-mediated phosphorylation of paxillin, thereby enhancing vinculin recruitment and FA maturation [11]. Early talin–vinculin pre-complexation can occur without tension, showing that force is not required for every step of adhesion formation [12]. Together, these findings provide a molecular basis for the modeled force-dependent maturation of FAs.

In the reciprocal direction, FA assembly influences AM tensile force through RhoA- and Rac1-related signaling cascades that exert opposing effects on force production. Force transmission through integrins can activate RhoA through the Rho guanine-nucleotide exchange factors LARG and GEF-H1, while adhesion-associated FAK/Pyk2 signaling can regulate RhoA through p190RhoGEF [13, 14]. Downstream RhoA–ROCK signaling promotes phosphorylation of the myosin regulatory light chain, inhibits myosin light-chain phosphatase, and supports AM contractility and traction generation [15–18]. Nascent or small adhesions recruit GIT1–*β*-PIX–PAK complexes that localize Rac1 activation and regulate adhesion and protrusion dynamics, while *β*-PIX can oppose myosin-II-dependent FA maturation [19– 21]. Although Rac1 can promote myosin II A recruitment in some cellular contexts [22], Rac1-mediated suppression of RhoA through the reactive-oxygen-species–p190RhoGAP pathway and PAK-mediated inhibition of myosin light-chain kinase provide molecular routes for an indirect inhibitory influence on AM tensile force [23–25]. Conversely, RhoA–ROCK signaling can suppress Rac through FilGAP [26]. Together, these pathways provide molecular routes for mutual inhibition between Rac1 and RhoA, consistent with their coordinated and antagonistic spatial dynamics and the capacity of this network to support bistable signaling behavior [27–29]. Accordingly, within the model, the RhoA- and Rac1-related pathways constitute positive and negative components, respectively, of the feedback from FA assembly to AM tensile force.

Each feedback interaction comprises multiple molecular and structural steps whose sequential progression generates a finite response latency. Because such multistep cascades can be coarse-grained as delayed input–output relations [30–32], we represent their combined temporal effect by a single effective feedback delay to retain this latency in a mathematically concise form. This effective delay is not assigned to a single biochemical reaction and does not assume that all underlying steps have the same duration.

### General delayed feedback function

Let *θ* denote dimensional time, and let *η* ≥ 0 denote a dimensional time delay. We consider regulation of *y*(*θ*) by the delayed value *x*(*θ* − *η*), where *x*(*θ*) *>* 0. We represent the regulatory dependence using the Hill function

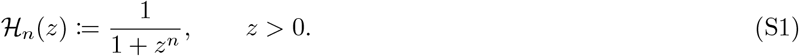

Here, *n* is a nonzero real-valued Hill exponent. As *z* increases, ℋ_*n*_(*z*) increases from zero to one for *n <* 0 and decreases from one to zero for *n >* 0, representing activating and decreasing regulatory dependences, respectively. The magnitude *n* controls the steepness of the response. Following explicit-delay formulations of biological regulatory networks [32], the corresponding delayed input to *y* is defined as

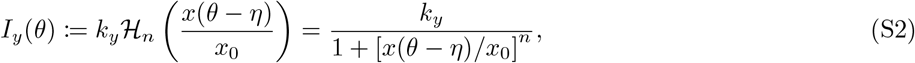

where *x*_0_ *>* 0 is the half-response scale and *k*_*y*_ *>* 0 is the saturating input magnitude.

### Dimensional mechanochemical model

Let *a*(*θ*) and *f* (*θ*) denote dimensional measures of FA assembly and AM tensile force, respectively. The FA assembly dynamics are written as

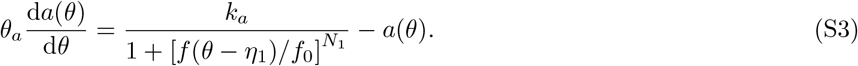

The exponent *N*_1_ *<* 0 makes the delayed Hill term an increasing function of *f* and represents a positive force-to-adhesion feedback in which AM tensile force promotes FA assembly and maturation. The scale *f*_0_ *>* 0 is the half-response force, *k*_*a*_ *>* 0 sets the saturating magnitude of the assembly input, *η*_1_ ≥ 0 denotes the dimensional time delay associated with the force-to-adhesion feedback, and *θ*_*a*_ *>* 0 sets the characteristic timescale over which the adhesion variable relaxes toward its input-dependent level. The linear loss term − *a*(*θ*) provides effective negative self-regulation of the adhesion variable by representing FA disassembly and turnover.

The dynamics of AM tensile force include two delayed adhesion-dependent inputs and are written as

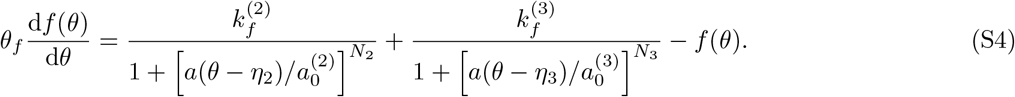

With *N*_2_ *<* 0 and *N*_3_ *>* 0, both Hill terms contribute positively to force production, but the RhoA-related term increases and the Rac1-related term decreases with FA assembly, generating positive and negative adhesion-to-force feedback, respectively. The parameters 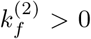 and 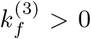 set the saturating magnitudes of the two inputs, and 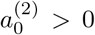 and 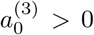 are their half-response adhesion scales. The parameters *η*_2_, *η*_3_ ≥ 0 denote the dimensional time delays associated with the RhoA- and Rac1-related feedback inputs, respectively. The parameter *θ*_*f*_ *>* 0 sets the characteristic timescale over which the force variable relaxes toward the level specified by the combined adhesion-dependent inputs. The linear loss term − *f* (*θ*) provides effective negative self-regulation of the force variable by representing relaxation of AM tensile force.

Because *a* and *f* are coarse-grained variables coupled within the same reciprocal feedback system, and no pronounced separation between their effective relaxation times is assumed, we use a common relaxation timescale *θ*_*R*_,

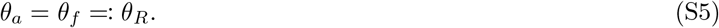

Motivated by the mutual inhibition between RhoA and Rac1, we coarse-grain their effects as opposing responses to the same FA assembly variable with a common half-response scale *a*_0_,

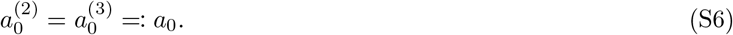

These assumptions define a symmetric coarse-grained model without asserting identical molecular rates or thresholds.

### Nondimensionalization

We define dimensionless time and state variables as

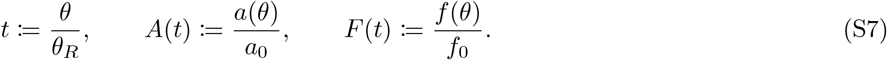

The dimensionless delays are defined by

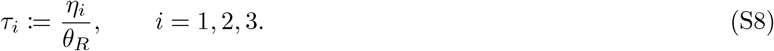

The delays *τ*_1_, *τ*_2_, and *τ*_3_ correspond to force-dependent adhesion assembly, RhoA-related positive feedback, and Rac1-related negative feedback, respectively. The dimensionless feedback strengths are defined by

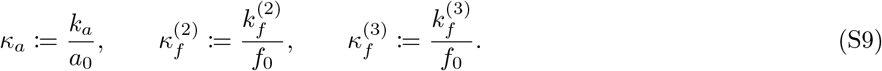

Substituting Eq. (S7)–Eq. (S9) into Eq. (S3) and Eq. (S4) gives

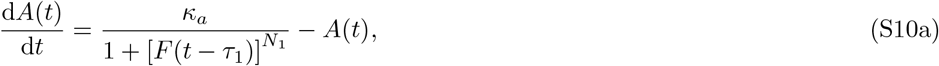

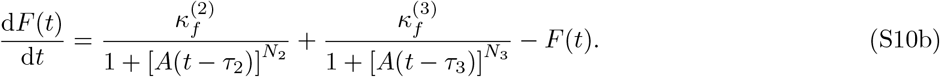

### Reduction to the three-parameter mechanochemical model

To isolate how feedback strength, feedback balance, and delay shape the system dynamics, we reduce the general nondimensional system to a symmetric three-parameter model. This reduction retains the positive force-to-adhesion feedback, the opposing RhoA- and Rac1-related adhesion-to-force feedbacks, and a finite feedback latency.

We set *N*_1_ = *N*_2_ = −2 for the two increasing Hill functions and *N*_3_ = 2 for the decreasing Hill function. The common exponent magnitude |*N*_*i*_ |= 2 is the smallest integer choice above unity and provides smooth sigmoidal responses with moderate nonlinearity without approaching an idealized step-function limit. These Hill exponents were fixed throughout the analysis and were not inferred from the experimental data as measures of molecular cooperativity.

To separate overall mechanochemical coupling from the relative balance of the two adhesion-to-force feedbacks, we set the force-to-adhesion strength equal to the total adhesion-to-force strength and define

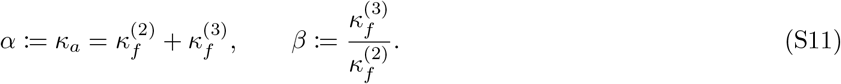

Here, *α* controls the common magnitude of the feedback inputs and thus the overall strength of coupling between FA assembly and AM tensile force. The ratio *β* quantifies the decreasing Rac1-related contribution relative to the increasing RhoA-related contribution and represents the balance between these opposing effects. Solving Eq. (S11) for the two adhesion-to-force feedback strengths gives

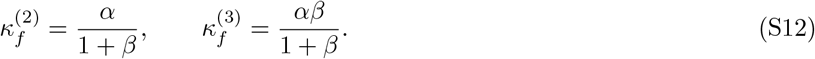

At fixed *α*, the increasing RhoA-related feedback strength dominates for *β <* 1, the two feedback strengths are equal at *β* = 1, and the decreasing Rac1-related feedback strength dominates for *β >* 1.

Direct measurement of each pathway-specific feedback delay is difficult because it integrates multiple molecular, cytoskeletal, and mechanical processes. We therefore represent the three delays by a common effective delay,

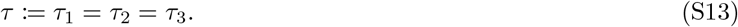

The dimensionless parameter *τ* summarizes the accumulated latency of signaling, cytoskeletal reorganization, force generation, and adhesion remodeling relative to the common relaxation timescale.

The resulting three-parameter dimensionless model used in the main text is given by

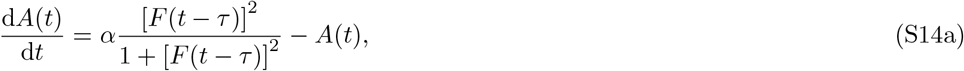

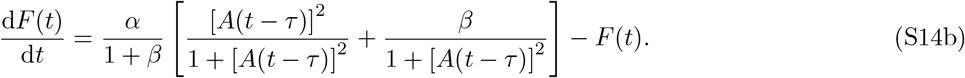

To make the role of *β* explicit, we define the delayed adhesion input as *A*_*τ*_ := *A*(*t* − *τ* ) *>* 0 and write the force equation as d*F/*d*t* = *G*_*F*_ (*A*_*τ*_ ; *α, β*) − *F* (*t*). The two Hill terms can then be combined as

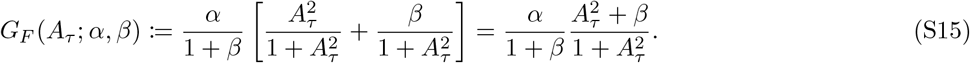

Its slope with respect to the delayed adhesion input is

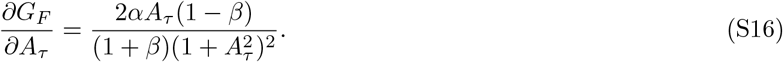

Because *α >* 0, *β >* 0, and *A*_*τ*_ *>* 0, the sign of this slope is determined by 1 − *β*. Thus, the net slope is positive for *β <* 1 (RhoA-dominant positive feedback), zero at *β* = 1 (*G*_*F*_ = *α/*2), and negative for *β >* 1 (Rac1-dominant negative feedback).

At a fixed point, *A*(*t* − *τ* ) = *A*(*t*) = *A*^∗^ and *F* (*t* − *τ* ) = *F* (*t*) = *F* ^∗^, so *τ* does not appear explicitly in the steady-state equations. The fixed-point locations are therefore independent of *τ*, although their stability and the time-dependent dynamics can depend on *τ* .

### State domain and history specification

We analyze the biologically relevant domain *A*(*t*) *>* 0 and *F* (*t*) *>* 0 with *α >* 0, *β >* 0, and *τ* ≥ 0. Unlike an ordinary differential equation, the delayed system at time *t* depends on its state at *t* − *τ*, so the values at *t* = 0 alone do not determine a solution. For *τ >* 0, we therefore specify positive history functions *A*(*t*) = *ϕ*_*A*_(*t*) and *F* (*t*) = *ϕ*_*F*_ (*t*) over *t* ∈ [− *τ*, 0] [33]. For *τ* = 0, the history specification reduces to the initial values *A*(0) *>* 0 and *F* (0) *>* 0. The histories and initial values used for each simulation are specified in the Materials and Methods and figure captions.

### Purpose and scope of the minimal mechanochemical model

The minimal model represents the collective effects of multiple molecular species, pathway-specific delays, response thresholds, and relaxation timescales using two state variables and three control parameters. The cited molecular mechanisms motivate the directions and signs of the effective feedback interactions but do not uniquely specify the functional forms or numerical parameter values. The model parameters were not fitted to the experimental RFP– vinculin or traction time series. Accordingly, the analysis addresses qualitative dynamical organization and does not aim to reproduce individual experimental time series quantitatively. The model was developed to investigate how a common mechanochemical feedback architecture can generate stable and oscillatory modes of cell–ECM adhesion dynamics. These modes offer a mechanistic interpretation of stable adhesion maintenance and dynamic adhesion remodeling, respectively.

## SUPPLEMENTARY TEXT S2. DERIVATION OF THE STABILITY AND BIFURCATION CONDITIONS

### Fixed points and parameterization

At a fixed point, *A*(*t* − *τ* ) = *A*(*t*) = *A*^∗^ and *F* (*t − τ* ) = *F* (*t*) = *F* ^∗^, so the delay drops out of the steady-state equations. Setting d*A/*d*t* = d*F/*d*t* = 0 in Eq. (S14) gives

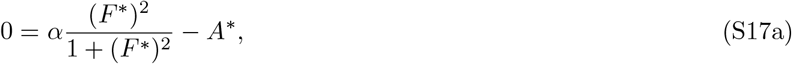

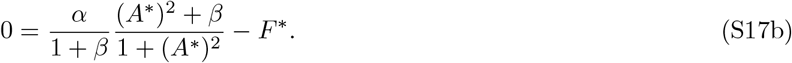

For prescribed (*α, β*), the positive solutions (*A*^∗^, *F* ^∗^) were determined numerically.

For the bifurcation derivations below, it is useful to parameterize *α* and *β* by the positive fixed-point coordinates (*A*^∗^, *F* ^∗^). Solving Eq. (S17a) for *α* gives

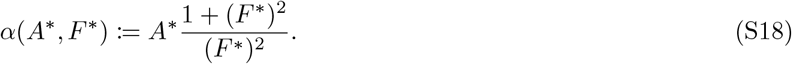

Solving Eq. (S17b) for *β* and substituting Eq. (S18) gives

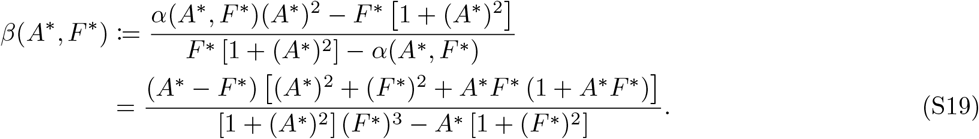

This parameterization is valid when the denominator in Eq. (S19) is nonzero and the resulting value satisfies *β >* 0. Within the positive fixed-point domain, Eq. (S19) is indeterminate only at (*A*^∗^, *F* ^∗^) = (1, 1), where the original fixed-point equations give *α* = 2 and are satisfied for every *β >* 0. This case was therefore handled separately whenever Eq. (S19) was used to parameterize fixed points or bifurcation points.

### Linearization and characteristic equation

To linearize Eq. (S14) around a positive fixed point (*A*^∗^, *F* ^∗^), we define the perturbations

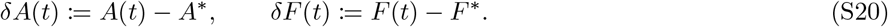

We also define the delayed states

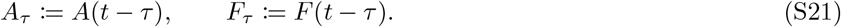

For *x >* 0, we define

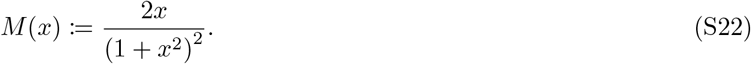

The derivatives of the force-to-adhesion and adhesion-to-force feedback functions at the fixed point are

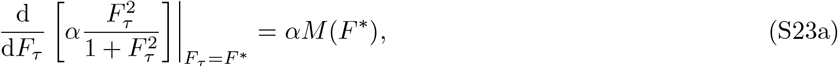

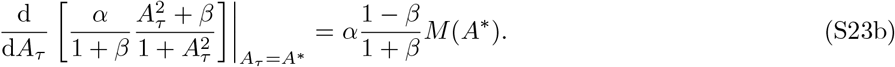

Retaining terms to first order in the perturbations gives

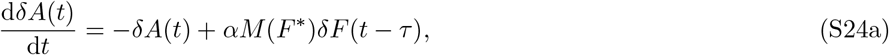

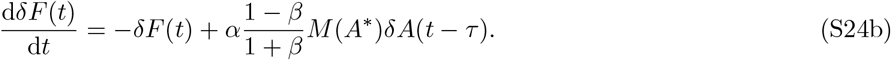

To derive the characteristic equation, we seek exponential solutions of the form

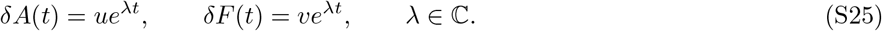

Substitution into Eq. (S24) gives

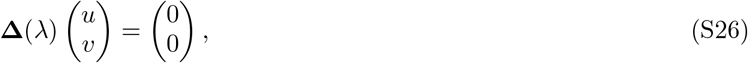

where the characteristic matrix is

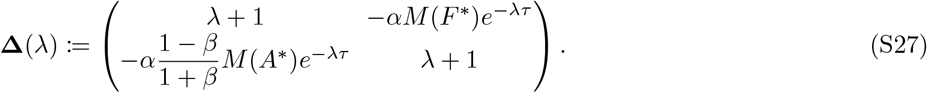

Nontrivial solutions require det **Δ**(*λ*) = 0.

For compactness, we define

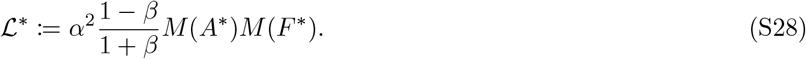

The characteristic function is then

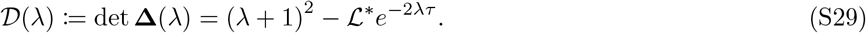

The characteristic roots satisfy

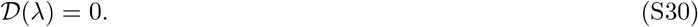

Because *M* (*A*^∗^) *>* 0 and *M* (*F* ^∗^) *>* 0, the sign of ℒ^∗^ is determined by 1 − *β*. Thus, ℒ^∗^ *>* 0 for *β <* 1, ℒ^∗^ = 0 for *β* = 1, and ^∗^ *<* 0 for *β >* 1. For *τ >* 0 and ℒ^∗^ /= 0, Eq. (S30) is transcendental and has infinitely many characteristic roots. The fixed point is linearly asymptotically stable if every characteristic root satisfies Re(*λ*) *<* 0.

### Saddle-node bifurcation condition

At a generic saddle-node bifurcation, two fixed-point branches meet at a nondegenerate fold, and the characteristic equation has a simple zero root. Substituting *λ* = 0 into Eq. (S30) gives

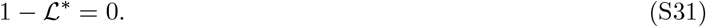

The zero-root condition is therefore

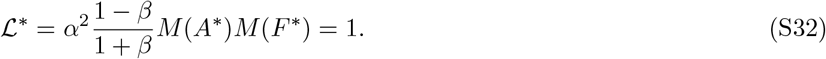

Because *α >* 0, *M* (*A*^∗^) *>* 0, and *M* (*F* ^∗^) *>* 0, this condition requires

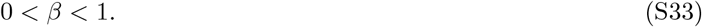

Neither the fixed-point equations nor Eq. (S32) contains *τ*, so the saddle-node boundary is independent of the feedback delay.

Differentiating Eq. (S29) with respect to *λ* and evaluating it at *λ* = 0 and ℒ^∗^ = 1 gives

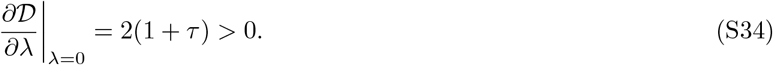

Because this derivative is nonzero for all *τ* ≥ 0, the zero characteristic root has algebraic multiplicity one, satisfying the simple-root condition required for a generic saddle-node bifurcation.

Using the fixed-point parameterization in Eq. (S18) and Eq. (S19), we define

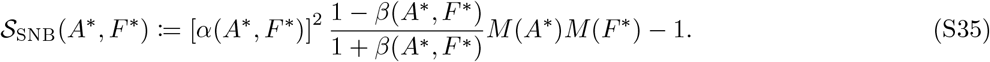

The parameterized fold locus was obtained by numerically identifying positive fixed-point coordinates satisfying

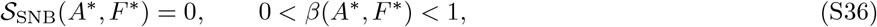

and mapping these coordinates to (*α, β*) using Eq. (S18) and Eq. (S19). Numerically computed fixed-point branches and nullcline analysis confirmed that the retained zero-root points were saddle-node folds at which two fixed-point branches met.

### Hopf bifurcation condition

A Hopf bifurcation requires a simple pair of complex-conjugate characteristic roots to cross the imaginary axis at *λ* = ±i*ω*, where *ω >* 0. Substituting *λ* = i*ω* into Eq. (S29) and separating the real and imaginary parts gives

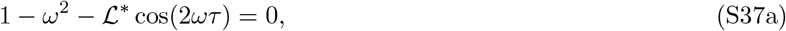

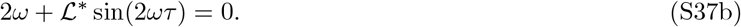

Squaring and adding these equations gives the magnitude condition

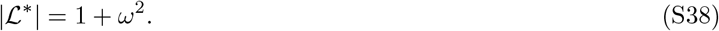

For *β <* 1, the linearized feedback gain satisfies ℒ^∗^ *>* 0. The formal purely imaginary solutions then satisfy

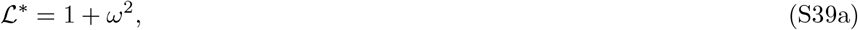

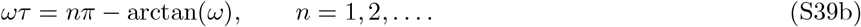

However, Eq. (S39a) requires ℒ^∗^ *>* 1. In this case, *D*(0) = 1 ^∗^ *<* 0, whereas *D*(*λ*) → + ∞ as the real variable *λ* →+ ∞.The characteristic equation therefore has a positive real root. Consequently, the fixed point is already unstable when the formal purely imaginary condition is satisfied, and this branch does not define the boundary at which a stable fixed point loses stability through a Hopf bifurcation.

For *β* = 1, ℒ^∗^ = 0, and the characteristic equation reduces to

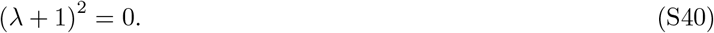

The only characteristic root is *λ* = −1, with algebraic multiplicity two, so no Hopf bifurcation occurs at *β* = 1. For *β >* 1, ℒ^∗^ *<* 0, and the purely imaginary roots satisfy

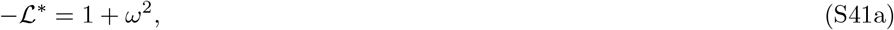

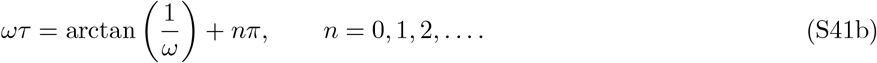

Equivalently, the phase condition satisfies

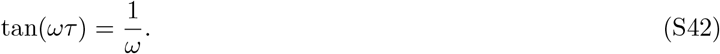

For a given fixed point at fixed (*α, β*) with *β >* 1, the characteristic roots at *τ* = 0 are

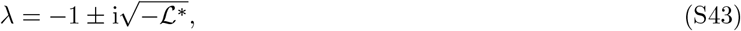

and therefore have negative real parts. For a fixed point satisfying −ℒ^∗^ *>* 1, Eq. (S41b) gives an increasing sequence of positive critical delays indexed by *n*. The *n* = 0 branch gives the smallest critical delay and hence the first loss of fixed-point stability, defining the linear Hopf boundary. Consequently, no Hopf bifurcation occurs in the nondelayed system (*τ* = 0).

Using the fixed-point parameterization in Eq. (S18) and Eq. (S19), we define

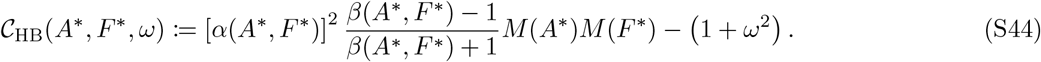

For each prescribed *τ >* 0, the first-branch frequency was determined from

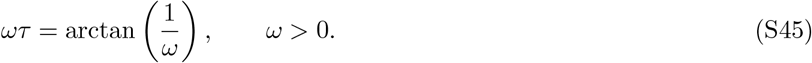

The corresponding Hopf boundary was obtained by identifying positive fixed-point coordinates satisfying

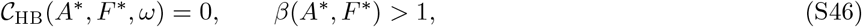

and mapping these coordinates to (*α, β*) using Eq. (S18) and Eq. (S19).

To verify that the purely imaginary roots are simple, we differentiate the characteristic function with respect to *λ*. Using *D*(i*ω*) = 0 gives

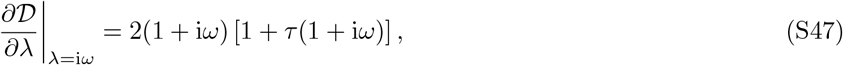

which is nonzero for *τ* ≥ 0 and *ω >* 0. Therefore, *λ* = ± i*ω* are simple characteristic roots. To determine the direction in which the characteristic roots cross the imaginary axis, we implicitly differentiate (*λ*) = 0 with respect to *τ* at fixed (*α, β*), giving

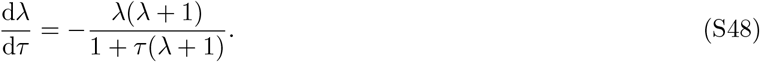

At *λ* = i*ω*, the real part satisfies

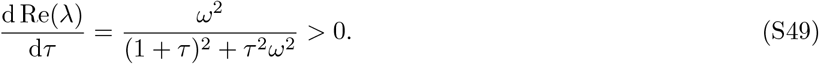

The conjugate pair therefore crosses the imaginary axis from the stable side to the unstable side at a nonzero rate as *τ* increases. This verifies the transversality condition and establishes the linear Hopf stability boundary. The linear analysis does not determine whether the bifurcation is supercritical or subcritical and does not establish periodic-orbit stability. Direct numerical integration nevertheless showed convergence to sustained periodic solutions at the examined parameter conditions on the oscillatory side of the boundary.

The corresponding phase relation between the critical linear perturbations can be obtained from the first row of Eq. (S26). At *λ* = i*ω*,

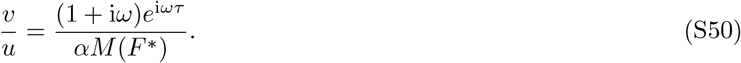

Because *αM* (*F* ^∗^) *>* 0, the phase of *F* relative to *A* is

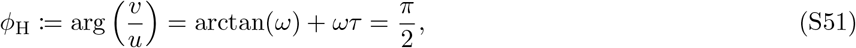

where the final equality follows from Eq. (S45). Thus, the critical linear mode at the first Hopf boundary has *F* leading *A* by one-quarter cycle. The phase relation of finite-amplitude periodic solutions was evaluated numerically.

## SUPPLEMENTARY MOVIE LEGENDS

### Supplementary Movie S1. Time-lapse visualization of RFP–vinculin fluorescence, cell-generated substrate wrinkles, and WFM-derived traction stress

Phase-contrast images, RFP–vinculin fluorescence images, and WFM-derived traction-stress vectors overlaid on the corresponding phase-contrast images are shown from left to right for recording R2, corresponding to Fig. 1A. Vector orientation indicates traction direction, whereas vector length and color indicate traction magnitude. Images were acquired sequentially at 2 min intervals, and the elapsed experimental time is indicated in the movie. Scale bar, 20 *µ*m.

## References

[1] K. Burridge, K. Fath, T. Kelly, G. Nuckolls, and C. Turner, Annual Review of Cell Biology 4, 487 (1988).

[2] B. Geiger, J. P. Spatz, and A. D. Bershadsky, Nature Reviews Molecular Cell Biology 10, 21 (2009).

[3] V. Vogel and M. Sheetz, Nature Reviews Molecular Cell Biology 7, 265 (2006).

[4] J. Z. Kechagia, J. Ivaska, and P. Roca-Cusachs, Nature Reviews Molecular Cell Biology 20, 457 (2019).

[5] N. Wang, J. P. Butler, and D. E. Ingber, Science 260, 1124 (1993).

[6] D. Choquet, D. P. Felsenfeld, and M. P. Sheetz, Cell 88, 39 (1997).

[7] R. J. Pelham, Jr. and Y.-L. Wang, Proceedings of the National Academy of Sciences of the United States of America 94, 13661 (1997).

[8] M. Chrzanowska-Wodnicka and K. Burridge, The Journal of Cell Biology 133, 1403 (1996).

[9] D. Riveline, E. Zamir, N. Q. Balaban, U. S. Schwarz, T. Ishizaki, S. Narumiya, Z. Kam, B. Geiger, and A. D. Bershadsky, The Journal of Cell Biology 153, 1175 (2001).

[10] C. Guilluy, V. Swaminathan, R. Garcia-Mata, E. T. O’Brien, R. Superfine, and K. Burridge, Nature Cell Biology 13, 722 (2011).

[11] S. L. Gupton and C. M. Waterman-Storer, Cell 125, 1361 (2006).

[12] A. del Rio, R. Perez-Jimenez, R. Liu, P. Roca-Cusachs, J. M. Fernandez, and M. P. Sheetz, Science 323, 638 (2009).

[13] P. Atherton, B. Stutchbury, D.-Y. Wang, D. Jethwa, R. Tsang, E. Meiler-Rodriguez, P. Wang, N. Bate, R. Zent, I. L. Barsukov, B. T. Goult, D. R. Critchley, and C. Ballestrem, Nature Communications 6, 10038 (2015).

[14] P. W. Oakes, E. Wagner, C. A. Brand, D. Probst, M. Linke, U. S. Schwarz, M. Glotzer, and M. L. Gardel, Nature Communications 8, 15817 (2017).

[15] K. Rottner, A. Hall, and J. V. Small, Current Biology 9, 640 (1999).

[16] A. S. Nimnual, L. J. Taylor, and D. Bar-Sagi, Nature Cell Biology 5, 236 (2003).

[17] J.-C. Kuo, X. Han, C.-T. Hsiao, J. R. Yates, and C. M. Waterman, Nature Cell Biology 13, 383 (2011).

[18] Y. Wu, K. Zhang, J. Seong, J. Fan, S. Chien, Y. Wang, and S. Lu, Scientific Reports 6, 29377 (2016).

[19] M. A. Fernández-Yaguë, E. N. Marquez, C. S. Poojari, J. Fu, Y. Wang, A. del Campo, and A. J. García, Science Advances 11, eadw6425 (2025).

[20] M. Beguerisse-Díaz, R. Desikan, and M. Barahona, Journal of The Royal Society Interface 13, 20160409 (2016).

[21] D. S. Glass, X. Jin, and I. H. Riedel-Kruse, Nature Communications 12, 1788 (2021).

[22] I. L. Novak, B. M. Slepchenko, A. Mogilner, and L. M. Loew, Physical Review Letters 93, 268109 (2004).

[23] V. S. Deshpande, M. Mrksich, R. M. McMeeking, and A. G. Evans, Journal of the Mechanics and Physics of Solids 56, 1484 (2008).

[24] E. S. Welf, H. E. Johnson, and J. M. Haugh, Molecular Biology of the Cell 24, 3945 (2013).

[25] A. Besser and U. S. Schwarz, Biophysical Journal 99, L10 (2010).

[26] C. E. Chan and D. J. Odde, Science 322, 1687 (2008).

[27] Z. Wu, S. V. Plotnikov, A. Y. Moalim, C. M. Waterman, and J. Liu, Biophysical Journal 112, 780 (2017).

[28] H. Li, D. Matsunaga, T. S. Matsui, H. Aosaki, G. Kinoshita, K. Inoue, A. Doostmohammadi, and S. Deguchi, Communications Biology 5, 361 (2022).

[29] C. D. Nobes and A. Hall, Cell 81, 53 (1995).

[30] J. L. MacKay and S. Kumar, Integrative Biology 6, 885 (2014).

[31] X.-D. Ren, W. B. Kiosses, D. J. Sieg, C. A. Otey, D. D. Schlaepfer, and M. A. Schwartz, Journal of Cell Science 113, 3673 (2000).

[32] B. D. Khalil, S. Hanna, B. A. Saykali, S. El-Sitt, A. Nasrallah, D. Marston, M. El-Sabban, K. M. Hahn, M. Symons, and M. El-Sibai, Experimental Cell Research 321, 109 (2014).

[33] A. Nayal, D. J. Webb, C. M. Brown, E. M. Schaefer, M. Vicente-Manzanares, and A. R. Horwitz, The Journal of Cell Biology 173, 587 (2006).

[34] N. Q. Balaban, U. S. Schwarz, D. Riveline, P. Goichberg, G. Tzur, I. Sabanay, D. Mahalu, S. Safran, A. Bershadsky, L. Addadi, and B. Geiger, Nature Cell Biology 3, 466 (2001).

[35] D. W. Dumbauld, H. Shin, N. D. Gallant, K. E. Michael, H. Radhakrishna, and A. J. García, Journal of Cellular Physiology 223, 746 (2010).

[36] J. Stricker, Y. Beckham, M. W. Davidson, and M. L. Gardel, PLOS ONE 8, e70652 (2013).

[37] L. Valon, A. Marín-Llaurado, T. Wyatt, G. Charras, and X. Trepat, Nature Communications 8, 14396 (2017).

[38] S. V. Plotnikov, A. M. Pasapera, B. Sabass, and C. M. Waterman, Cell 151, 1513 (2012).

[39] K. Sao, T. M. Jones, A. D. Doyle, D. Maity, G. Schevzov, Y. Chen, P. W. Gunning, and R. J. Petrie, Molecular Biology of the Cell 30, 1170 (2019).

[40] E. A. Cavalcanti-Adam, T. Volberg, A. Micoulet, H. Kessler, B. Geiger, and J. P. Spatz, Biophysical Journal 92, 2964 (2007).

[41] H. B. Schiller, M.-R. Hermann, J. Polleux, T. Vignaud, S. Zanivan, C. C. Friedel, Z. Sun, A. Raducanu, K.-E. Gottschalk, M. Théry, M. Mann, and R. Fassler, Nature Cell Biology 15, 625 (2013).

[42] C. Grashoff, B. D. Hoffman, M. D. Brenner, R. Zhou, M. Parsons, M. T. Yang, M. A. McLean, S. G. Sligar, C. S. Chen, T. Ha, and M. A. Schwartz, Nature 466, 263 (2010).

[43] A. M. Pasapera, I. C. Schneider, E. Rericha, D. D. Schlaepfer, and C. M. Waterman, The Journal of Cell Biology 188, 877 (2010).

[44] P. W. Oakes, Y. Beckham, J. Stricker, and M. L. Gardel, The Journal of Cell Biology 196, 363 (2012).

[45] B. Hinz, V. Dugina, C. Ballestrem, B. Wehrle-Haller, and C. Chaponnier, Molecular Biology of the Cell 14, 2508 (2003).

[46] S. S. Chang, A. D. Rape, S. A. Wong, W.-H. Guo, and Y.-L. Wang, Molecular Biology of the Cell 30, 3104 (2019).

[47] H. Xu, S. Donegan, J. M. Dreher, A. J. Stark, E. P. Canović, D. Stamenović, and M. L. Smith, Acta Biomaterialia 113, 372 (2020).

[48] J. Stricker, Y. Aratyn-Schaus, P. W. Oakes, and M. L. Gardel, Biophysical Journal 100, 2883 (2011).

[49] R. Pankov, E. Cukierman, B.-Z. Katz, K. Matsumoto, D. C. Lin, S. Lin, C. Hahn, and K. M. Yamada, The Journal of Cell Biology 148, 1075 (2000).

[50] B.-Z. Katz, E. Zamir, A. Bershadsky, Z. Kam, K. M. Yamada, and B. Geiger, Molecular Biology of the Cell 11, 1047 (2000).

[51] E. Matsumoto and S. Deguchi, Journal of Theoretical Biology 634, 112571 (2026).

[52] J. Fouchard, C. Bimbard, N. Bufi, P. Durand-Smet, A. Proag, A. Richert, O. Cardoso, and A. Asnacios, Proceedings of the National Academy of Sciences of the United States of America 111, 13075 (2014).

[53] V. F. Fiore, P. W. Strane, A. V. Bryksin, E. S. White, J. S. Hagood, and T. H. Barker, The Journal of Cell Biology 211, 173 (2015).

[54] X.-D. Ren, W. B. Kiosses, and M. A. Schwartz, The EMBO Journal 18, 578 (1999).

[55] A. Y. Mitrophanov and E. A. Groisman, BioEssays 30, 542 (2008).

[56] J. M. Whitacre, Frontiers in Genetics 3, 67 (2012).

[57] D. Mitrossilis, J. Fouchard, D. Pereira, F. Postic, A. Richert, M. Saint-Jean, and A. Asnacios, Proceedings of the National Academy of Sciences of the United States of America 107, 16518 (2010).

[58] A. M. Rosales, S. L. Vega, F. W. DelRio, J. A. Burdick, and K. S. Anseth, Angewandte Chemie International Edition 56, 12132 (2017).

[59] P. Linke, N. Munding, E. Kimmle, S. Kaufmann, K. Hayashi, M. Nakahata, Y. Takashima, M. Sano, M. Bastmeyer, T. Holstein, S. Dietrich, C. Müller-Tidow, A. Harada, A. D. Ho, and M. Tanaka, Advanced Healthcare Materials 13, e2302607 (2024).

[60] Z. Zhang, H. Zhu, G. Zhao, Y. Miao, L. Zhao, J. Feng, H. Zhang, R. Miao, L. Sun, B. Gao, W. Zhang, Z. Wang, J. Zhang, Y. Zhang, H. Guo, F. Xu, T. J. Lu, G. M. Genin, and M. Lin, Advanced Science 10, e2302421 (2023).

[61] M. Maraldi, C. Valero, and K. Garikipati, Biophysical Journal 106, 1890 (2014).

[62] X. Cao, Y. Lin, T. P. Driscoll, J. Franco-Barraza, E. Cukierman, R. L. Mauck, and V. B. Shenoy, Biophysical Journal 109, 1807 (2015).

[63] G. R. McNicol, M. J. Dalby, and P. S. Stewart, Journal of Theoretical Biology 596, 111965 (2025).

[64] M. Cirit, M. Krajcovic, C. K. Choi, E. S. Welf, A. F. Horwitz, and J. M. Haugh, PLoS Computational Biology 6, e1000688 (2010).

[65] S. Talwar, A. Kant, T. Xu, V. B. Shenoy, and R. K. Assoian, Cell Reports 35, 109019 (2021).

[66] T. Erdmann and U. S. Schwarz, Biophysical Journal 91, L60 (2006).

[67] P. Liu, Q. Wang, X. Dai, L. Pei, J. Wang, W. Zhao, H. E. Johnson, M. Yao, and A. K. Efremov, Nature Physics 21, 1431 (2025).

[68] D. Ghosh, S. Ghosh, and A. Chaudhuri, Biophysical Journal 121, 1753 (2022).

[69] V. Andasari and S. Syafiie, bioRxiv 10.1101/2025.05.06.652413 (2025), preprint; posted May 8, 2025.

[70] S. Deguchi, Y. Nagasawa, A. C. Saito, T. S. Matsui, S. Yokoyama, and M. Sato, Biotechnology Letters 36, 507 (2014).

[71] H. Li, D. Matsunaga, T. S. Matsui, H. Aosaki, and S. Deguchi, Biochemical and Biophysical Research Communications 530, 527 (2020).

[72] K. Burridge and C. Guilluy, Experimental Cell Research 343, 14 (2016).

[73] J. P. ten Klooster, Z. M. Jaffer, J. Chernoff, and P. L. Hordijk, The Journal of Cell Biology 172, 759 (2006).

[74] Y. Lim, S.-T. Lim, A. Tomar, M. Gardel, J. A. Bernard-Trifilo, X. L. Chen, S. A. Uryu, R. Canete-Soler, J. Zhai, H. Lin, W. W. Schlaepfer, P. Nalbant, G. Bokoch, D. Ilic, C. Waterman-Storer, and D. D. Schlaepfer, The Journal of Cell Biology 180, 187 (2008).

[75] Y. Ohta, J. H. Hartwig, and T. P. Stossel, Nature Cell Biology 8, 803 (2006).

[76] M. Machacek, L. Hodgson, C. Welch, H. Elliott, O. Pertz, P. Nalbant, A. Abell, G. L. Johnson, K. M. Hahn, and G. Danuser, Nature 461, 99 (2009).

[77] K. M. Byrne, N. Monsefi, J. C. Dawson, A. Degasperi, J.-C. Bukowski-Wills, N. Volinsky, M. Dobrzyński, M. R. Birtwistle, M. A. Tsyganov, A. Kiyatkin, K. Kida, A. J. Finch, N. O. Carragher, W. Kolch, L. K. Nguyen, A. von Kriegsheim, and B. N. Kholodenko, Cell Systems 2, 38 (2016).

[78] K. Kimura, M. Ito, M. Amano, K. Chihara, Y. Fukata, M. Nakafuku, B. Yamamori, J. Feng, T. Nakano, K. Okawa, A. Iwamatsu, and K. Kaibuchi, Science 273, 245 (1996).

[79] L. C. Sanders, F. Matsumura, G. M. Bokoch, and P. de Lanerolle, Science 283, 2083 (1999).

[80] G. Totsukawa, Y. Yamakita, S. Yamashiro, D. J. Hartshorne, Y. Sasaki, and F. Matsumura, The Journal of Cell Biology 150, 797 (2000).

[81] J. Srividhya, M. S. Gopinathan, and S. Schnell, Biophysical Chemistry 125, 286 (2007).

[82] T. Erneux, Applied Delay Differential Equations, Surveys and Tutorials in the Applied Mathematical Sciences, Vol. 3 (Springer, New York, NY, 2009).

## SUPPLEMENTARY REFERENCES

[1] V. Vogel and M. Sheetz, Nature Reviews Molecular Cell Biology 7, 265 (2006).

[3] K. Burridge and C. Guilluy, Experimental Cell Research 343, 14 (2016).

[4] N. Q. Balaban, U. S. Schwarz, D. Riveline, P. Goichberg, G. Tzur, I. Sabanay, D. Mahalu, S. Safran, A. Bershadsky, L. Addadi, and B. Geiger, Nature Cell Biology 3, 466 (2001).

[5] D. Riveline, E. Zamir, N. Q. Balaban, U. S. Schwarz, T. Ishizaki, S. Narumiya, Z. Kam, B. Geiger, and A. D. Bershadsky, The Journal of Cell Biology 153, 1175 (2001).

[6] A. del Rio, R. Perez-Jimenez, R. Liu, P. Roca-Cusachs, J. M. Fernandez, and M. P. Sheetz, Science 323, 638 (2009).

[7] C. Grashoff, B. D. Hoffman, M. D. Brenner, R. Zhou, M. Parsons, M. T. Yang, M. A. McLean, S. G. Sligar, C. S. Chen, T. Ha, and M. A. Schwartz, Nature 466, 263 (2010).

[8] P. Atherton, B. Stutchbury, D.-Y. Wang, D. Jethwa, R. Tsang, E. Meiler-Rodriguez, P. Wang, N. Bate, R. Zent, I. L. Barsukov, B. T. Goult, D. R. Critchley, and C. Ballestrem, Nature Communications 6, 10038 (2015).

[9] M. Yao, B. T. Goult, B. Klapholz, X. Hu, C. P. Toseland, Y. Guo, P. Cong, M. P. Sheetz, and J. Yan, Nature Communications 7, 11966 (2016).

[10] Y. Wang, M. Yao, K. B. Baker, R. E. Gough, S. Le, B. T. Goult, and J. Yan, Journal of the American Chemical Society 143, 14726 (2021).

[11] A. M. Pasapera, I. C. Schneider, E. Rericha, D. D. Schlaepfer, and C. M. Waterman, The Journal of Cell Biology 188, 877 (2010).

[12] S. J. Han, E. V. Azarova, A. J. Whitewood, A. Bachir, E. Guttierrez, A. Groisman, A. R. Horwitz, B. T. Goult, K. M. Dean, and G. Danuser, eLife 10, e66151 (2021).

[13] Y. Lim, S.-T. Lim, A. Tomar, M. Gardel, J. A. Bernard-Trifilo, X. L. Chen, S. A. Uryu, R. Canete-Soler, J. Zhai, H. Lin, W. W. Schlaepfer, P. Nalbant, G. Bokoch, D. Ilic, C. Waterman-Storer, and D. D. Schlaepfer, The Journal of Cell Biology 180, 187 (2008).

[14] C. Guilluy, V. Swaminathan, R. Garcia-Mata, E. T. O’Brien, R. Superfine, and K. Burridge, Nature Cell Biology 13, 722 (2011).

[15] M. Chrzanowska-Wodnicka and K. Burridge, The Journal of Cell Biology 133, 1403 (1996).

[16] K. Kimura, M. Ito, M. Amano, K. Chihara, Y. Fukata, M. Nakafuku, B. Yamamori, J. Feng, T. Nakano, K. Okawa, A. Iwamatsu, and K. Kaibuchi, Science 273, 245 (1996).

[17] G. Totsukawa, Y. Yamakita, S. Yamashiro, D. J. Hartshorne, Y. Sasaki, and F. Matsumura, The Journal of Cell Biology 150, 797 (2000).

[18] P. W. Oakes, E. Wagner, C. A. Brand, D. Probst, M. Linke, U. S. Schwarz, M. Glotzer, and M. L. Gardel, Nature Communications 8, 15817 (2017).

[19] A. Nayal, D. J. Webb, C. M. Brown, E. M. Schaefer, M. Vicente-Manzanares, and A. R. Horwitz, The Journal of Cell Biology 173, 587 (2006).

[20] J. P. ten Klooster, Z. M. Jaffer, J. Chernoff, and P. L. Hordijk, The Journal of Cell Biology 172, 759 (2006).

[21] J.-C. Kuo, X. Han, C.-T. Hsiao, J. R. Yates, and C. M. Waterman, Nature Cell Biology 13, 383 (2011).

[22] A. M. Pasapera, S. V. Plotnikov, R. S. Fischer, L. B. Case, T. T. Egelhoff, and C. M. Waterman, Current Biology 25, 175 (2015).

[23] E. E. Sander, J. P. ten Klooster, S. van Delft, R. A. van der Kammen, and J. G. Collard, The Journal of Cell Biology 147, 1009 (1999).

[24] L. C. Sanders, F. Matsumura, G. M. Bokoch, and P. de Lanerolle, Science 283, 2083 (1999).

[25] A. S. Nimnual, L. J. Taylor, and D. Bar-Sagi, Nature Cell Biology 5, 236 (2003).

[26] Y. Ohta, J. H. Hartwig, and T. P. Stossel, Nature Cell Biology 8, 803 (2006).

[27] K. Rottner, A. Hall, and J. V. Small, Current Biology 9, 640 (1999).

[28] M. Machacek, L. Hodgson, C. Welch, H. Elliott, O. Pertz, P. Nalbant, A. Abell, G. L. Johnson, K. M. Hahn, and G. Danuser, Nature 461, 99 (2009).

[29] K. M. Byrne, N. Monsefi, J. C. Dawson, A. Degasperi, J.-C. Bukowski-Wills, N. Volinsky, M. Dobrzyński, M. R. Birtwistle, M. A. Tsyganov, A. Kiyatkin, K. Kida, A. J. Finch, N. O. Carragher, W. Kolch, L. K. Nguyen, A. von Kriegsheim, and B. N. Kholodenko, Cell Systems 2, 38 (2016).

[30] J. Srividhya, M. S. Gopinathan, and S. Schnell, Biophysical Chemistry 125, 286 (2007).

[31] M. Beguerisse-Díaz, R. Desikan, and M. Barahona, Journal of The Royal Society Interface 13, 20160409 (2016).

[32] D. S. Glass, X. Jin, and I. H. Riedel-Kruse, Nature Communications 12, 1788 (2021).

[33] T. Erneux, Applied Delay Differential Equations, Surveys and Tutorials in the Applied Mathematical Sciences, Vol. 3 (Springer, New York, NY, 2009).

